# A systematic screen of breast cancer patients’ exomes for retrotransposon insertions reveals disease associated genes

**DOI:** 10.1101/2020.06.04.123240

**Authors:** Sylvia De Brakeleer, Jacques De Grève, Erik Teugels

## Abstract

**Background:** Retrotransposons are genetic elements that jump within the genome via an RNA intermediate. Although they had a strong impact on human genome evolution, only a very tiny fraction of them can be reactivated nowadays, most often with neutral or detrimental consequences. The pathological outcomes associated with such genetic alterations are poorly investigated in the clinic, merely due to their difficult detection.

**Results:** We developed a strategy to detect rare retrotransposon mediated insertions in Whole Exome Sequencing data from 65 familial breast cancer patients. When restricting our search to high confidence retrotransposition events occurring in less than 10% of the samples, we identified only ten different Alu elements, two L1 elements, one SVA and two processed pseudogenes. Only two of these insertions occurred within protein coding sequences and interestingly, several of the targeted genes have been previously linked to cancer, in three cases even to increased breast cancer risk (*GHR, DMBT1* and *NEK10*). When investigating the molecular consequences of four Alu insertions at the mRNA level, we found that the element present in the 3’UTR of *GHR* repressed expression of the corresponding allele. oMreover, the analysis of a near exonic Alu insertion in *PTPN14* (a mediator of *P53* tumor suppressor activity) revealed that this gene was imprinted and that the presence of an intronic Alu element can lead to loss of imprinting.

**Conclusions:** Our data underline the relevance of incorporating the search for uncommon retrotransposition events in Next Generation Sequencing pipelines when analyzing patients with a suspected genetic disease.

## BACKGROUND

More than 50% of the human genome is built up with sequences that originated from the activity of transposable elements[1], mainly retrotransposons which are able to move to new locations in the genome via an RNA intermediate using the copy/paste principle[2-4]. L1 retrotransposons belong to the LINE (long interspersed element) superfamily and are the only transposons still active in human. Full length L1 elements (∼6kb long) code for the proteins necessary for retrotransposition, including a reverse transcriptase. Although L1 sequences represent about 17% of the human genome, only a very small fraction of them (80-100 copies) retained their transposition capacity[5, 6]. Alu and SVA retrotransposons (about 300 and 1400bp long respectively) are SINEs (short interspersed elements) and do not code for proteins. Rather, they are fully dependent on L1 retrotransposition and parasitize its propagation system. Accidentally, cellular RNA can also misuse the L1 retrotransposition machinery resulting in the generation of processed pseudogenes[7]. The CpG rich sequences present in L1 promoters, Alu and SVA elements are sites for DNA methylation and heterochromatin formation, causing epigenetic silencing[8]. Interestingly, the vast majority of evolutionary stabilized Alu insertions are located in gene-rich regions. They are often embedded in sequences encoding pre-mRNAs or mature mRNAs, usually as part of their introns or UTRs were they can potentially contribute to transcriptome variation[9].

In 1985 the first de novo Alu element insertion was reported in a B cell lymphoma[10]. Three years later, Kazazian et al.[11] reported a Haemophilia A causing L1 element insertion in germline DNA. In a review article published in 2016[6], the list of retrotransposons associated with human diseases counts 124 entries. Obviously, the number of disease causing retrotransposition events identified in human[12] is increasing steadily, but the attention these mutations receive in the clinic is still very poor when compared to classical mutational events (nucleotide substitutions and small insertions/deletions). When searching for breast cancer (BC) predisposing mutations in high risk families, we previously identified one Alu insertion in exon 11 of *BRCA1* and one Alu insertion in exon 3 of the *BRCA2* gene[13], this last one appeared later on to be a recurrent founder mutation restricted to the Portuguese population[14]. When Next Generation Sequencing tools were introduced in the diagnostic field, only few labs adapted their IN SILICO pipeline to allow the identification of pathogenic retrotransposition events in the genes they analyze[15, 16]. In the present work, we developed a strategy allowing the detection of uncommon retrotransposition events in Whole Exome Sequencing (WES) data using BAM files generated from a cohort of familial breast cancer patients previously used to identify more classical cancer predisposing mutations[17]. In a second step, we investigated the molecular consequences generated by 5 candidate pathogenic Alu insertions revealed during our screen, and highlight the relevance of screening for rare retrotransposition events in the clinical context.

## RESULTS

### IN SILICO screen for retrotransposition mediated insertions

Because of their shared mechanisms of transposition using an RNA molecule as intermediate, retrotransposons all bear a long polyA tail at their 3’ extremity. Also, their specific mechanism for insertion into the target DNA results in the duplication of a short sequence at the position where the retrotransposon has integrated. This duplicated sequence (called Target Site Duplication or TSD) is usually between 4 and 20 nucleotides long and flanks the retrotransposon specific sequences. We exploited those two characteristics to identify retrotransposon insertions not present in the human reference genome (hg19) using WES data generated with the SeqCap EZ Exome v3.0 kit from Roche. Paired-end sequencing usually generates read pairs that perfectly map in close proximity on a reference genome. However, the presence of a non-reference retrotransposon insertion will result in read pairs that do not map closely (Figure1). We selected these discordantly mapped read pairs by applying the open-source RetroSeq software[18] on the BAMfiles previously obtained from 65 BC patients[17] (positive *BRCA1* control included), with a first additional restriction that one read of the discordantly mapped read pairs must harbor a long polyA stretch. On average, we obtained 987 candidate insertions for each patient. When introducing additional filtering steps (using an Excel program) to remove false positives and frequently occurring non reference polymorphic insertions, we ended with a list counting only 32 different candidate polyA containing insertion sites for the pooled 65 patients (Additional file 1). A detailed description of our detection strategy and its application on a positive control sample is presented in the eMthod section, wherein we also compare two widely used exome enrichment kits.

**Figure 1:**
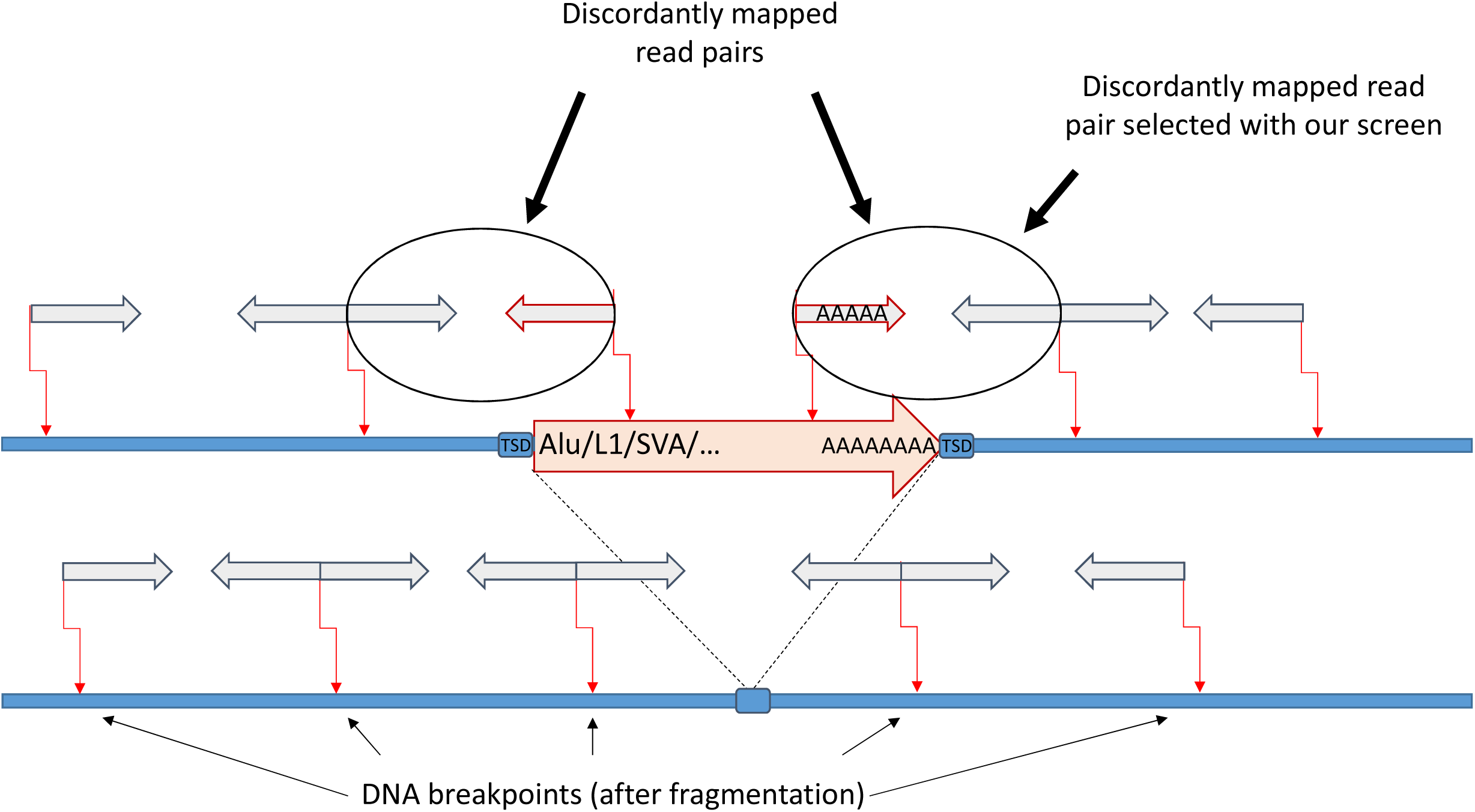
Title: Schematic representation of the “discordantly mapped read pairs” used for the detection of polymorphic retrotransposons. Legend: After genome fragmentation (broken arrows), size selected DNA fragments are sequenced from both ends (paired-end sequencing). DNA fragments having one breakpoint in a gene and the other breakpoint in a retrotransposon not included in the reference human genome will generate read pairs (grey filled arrows) that do not map at nearby positions on the reference genome after alignment. The discordantly mapped read pairs obtained by analysis of the full exome of a patient (traceable in the corresponding BAMfile) are further selected for the presence of a long polyA stretch in one of the mate reads. An additional landmark for retrotransposition is the presence of a target site duplication (TSD) flanking the transposon.

### High confident retrotransposition mediated insertions

When performing a manual selection (using the Integrative Genomics Viewer: IGV[19]) for the presence of a TSD on the 32 polyA containing insertions we had identified, we ended up with 18 high confidence candidates retrotransposon insertions (Table 1). The further characterization of these retrotransposition mediated insertions (12 Alu, 2 L1, 1 SVA and 3 pseudogenes, see Table 1) was made possible by the presence of a stretch of nucleotides (maximum ∼50bp long) in a fraction of the aligned read pairs that do not match with the reference sequence, but corresponded the 5’ extremity of the inserted sequence. These sequences could be identified using available web tools (see eMthods). A representative screenshot showing the outcome of an IGV analysis of the genomic region wherein the Alu element integrated into the BRCA1 gene of our positive control sample is presented in Figure 2. Only 2 out of the 18 retrotransposon mediated events occurred within protein coding regions (Alu element in *ZNF442* and *UQCR10* transcript in *C1orf194*, see Table 1) which might be surprising since WES probes are expected to be specifically designed towards exonic sequences. Indeed, according to the kit provider (Roche) only protein coding parts of the transcripts are targeted, but for exons that are smaller than 100 bp, the target region is extended to 100 bp. Nevertheless, we expect that the large majority of retrotransposition mediated insertions occurring in the well covered protein coding regions will be picked up with our screen, their low incidence being in agreement with the literature[5, 20, 21]. Conversely, we are convinced that a significant fraction of the polymorphic retrotransposons located in the vicinity of protein coding regions (regions often poorly or not covered at all by WES) is missed with our WES based approach, although the large majority of identified insertions (15/17, positive control excluded) are located in intronic, 3’UTR, promoter or even intergenic sequences. As a sufficient coverage at sequencing is essential to allow the identification of polymorphic retrotransposons in the non-coding regions, we wanted to estimate the number of false negative carriers of such highly confident retrotransposition mediated insertions among the 65 samples we investigated. Therefore, all samples were rescreened for the presence of the 18 IN SILICO identified insertion types using IGV (see Table 1). All insertions identified with the IN SILICO pipeline could be confirmed with IGV analysis (no false positives). As expected, IGV inspection revealed higher rates of carriers for 9 insertions (Table 1). In two cases, insertion carriership was even observed in 50% of the samples, with IGV and IN SILICO data discrepancies being well explainable by the low coverage at sequencing. Of note, about half of the high confident retrotransposon insertions we identified are reported in the database of retrotransposon insertion polymorphisms in human (dbRIP, see Table 1).

**Table 1:**
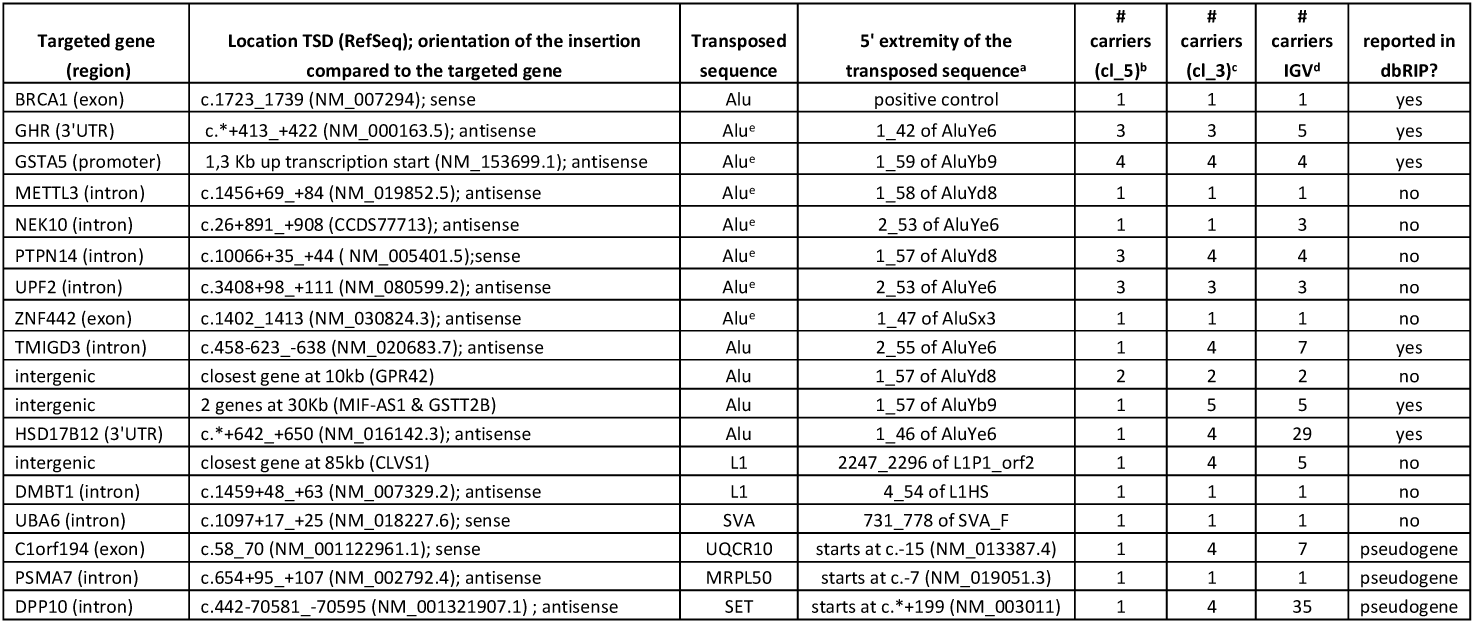
List with the 18 high confidence retrotransposition mediated insertions detected in 65 familial BC patients

**Figure 2:**
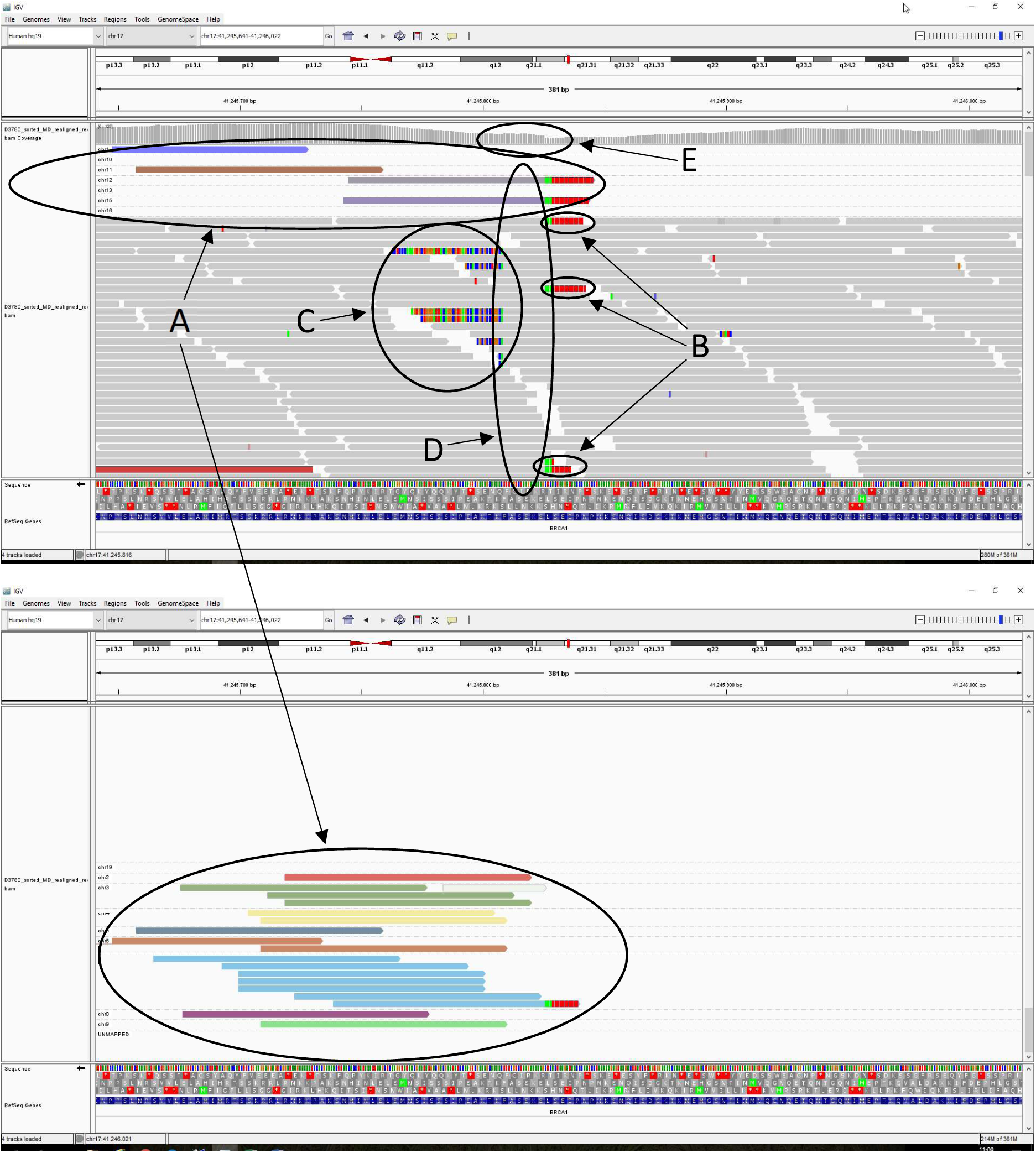
Title: Screenshots of IGV outputs visualizing the Alu element insertion in the positive control sample. Legend: The screenshots are restricted to the genomic region where the Alu element insertion occurred (c.1739_1740insAlu in *BRCA1*) in the positive control sample. The BAMfile used for IGV visualization was generated from WES data obtained with the exome capture kit SeqCap EZ Exome v3.0 from Roche. Group alignment is by chromosome of mate. Note the presence of a high number of reads (22) whose mate reads are located on another chromosome (encircled and labelled “A”). All those BRCA1 specific reads point in the same direction, the 3’ extremity of the Alu insertion that ends with a long polyA stretch (3 of these 22 reads already end with a small polyA stretch). Consequently, the mates of these 22 reads have a high probability to contain a long polyA stretch and to be picked up with our detection strategy. Five such discordantly mapped read pairs are sufficient to retain this particular genomic region as a candidate insertion site. Note also that a number of correctly mapped read pairs (labelled “B” and “C”) start with a sequence that do not correspond to the *BRCA1* sequence. The reads labelled “B” confirm the presence (and position) of a polyA stretch adjacent to the *BRCA1* sequence, while the reads labelled “C” allow the identification of the inserted sequence most upstream of the polyA stretch, in this case an Alu element. Inserted retrotransposon sequences can be identified making use of the Dfam [58] database (https://dfam.org/home), processed pseudogenes with BLAST (https://blast.ncbi.nlm.nih.gov/Blast.cgi). Finally, and characteristic for retrotransposition events, the inserted element is flanked by a short repeated sequence (TSD, 17bp long in this case) as is indicated with label “D”. Often, a sharp increase of the coverage at the level of the TSD is observed (less pronounced in this example, see label “E”).

### Retrotransposon insertions in non-coding regions can affect allelic expression

To investigate the potential consequences of Alu insertions located in non-coding gene regions at the molecular level[22], we collected blood samples from available carriers (and non-carriers for control) and extracted the corresponding RNA material. Since allele specific expression levels can be monitored by Sanger sequencing only when the Alu carrier is also heterozygous for a coding SNP in the same gene, the number of suitable insAlu carriers among the 65 investigated patients was not always sufficient and additional familial BC patients were genotyped using mutation specific hemi-nested PCRs (Table 2). Sufficient blood samples could be collected to initiate the analysis of 5 Alu targeted genes (*GHR, GSTA5, NEK10, PTPN14* and *UPF2*). Sequence analyses of the cDNA regions flanking the allele discriminating SNPs revealed that the intronic Alu insertions in *UPF2* and *NEK10* do not result in an obvious decrease or increase of the mRNA levels generated from the retrotransposon targeted allele (Figure 3). In sharp contrast, the Alu insertion in the 3’UTR of *GHR* clearly resulted in allele silencing (Figure 4A) suggesting that this Alu sequence may contain sites for microRNA directed degradation, a mechanism previously proposed to explain the occurrence of evolutionary stabilized Alu derived microRNA binding sites in the 3’UTRs of specific gene[23]. Conversely, the two available carriers of the intronic Alu insertion in *PTPN14* seem to express both alleles equally well but among the three non-carriers, one expresses virtually only one allele while the two others show a reduced expression of this same allele (Figure 4B). Partial or near absolute mono-allelic expression of *PTPN14* is in agreement with a study listing this gene as a high-confidence imprinted human gene candidate with maternal expression[24]. On the other hand, the apparent loss of imprinting (LOI) observed in the Alu carriers suggests that the integration of one single Alu element into the gene sequence of the imprinted allele would be sufficient to fully reactivate this allele, which matches with earlier observations reporting a sharp inflection in SINE content at transitions from imprinted to non-imprinted genomic regions[25]. Unfortunately, differential expression of the *GSTA5* Alu containing allele (insertion occurred in the promoter region) could not be investigated due to the very low expression level of *GSTA5* in leucocytes compared to *GSTA1* and *GSTA2* (see GeneCards.org), three homologues that are hard to discriminate when using PCR based assays (only GSTA1 and/or GSTA2 sequences could be detected after Sanger sequencing even by using gene discriminating primers for PCR, results not shown).

**Table 2:** Results from insAlu genotyping experiments performed on 710 non related familial BC patients

| Alu targeted gene | # carriers out of 710 |
| --- | --- |
| UPF2 | 107 |
| PTPN14 | 10 |
| GHR | 17 |
| ZNF442 | 1 |
| METTL3 | 1 |

**Figure 3:**
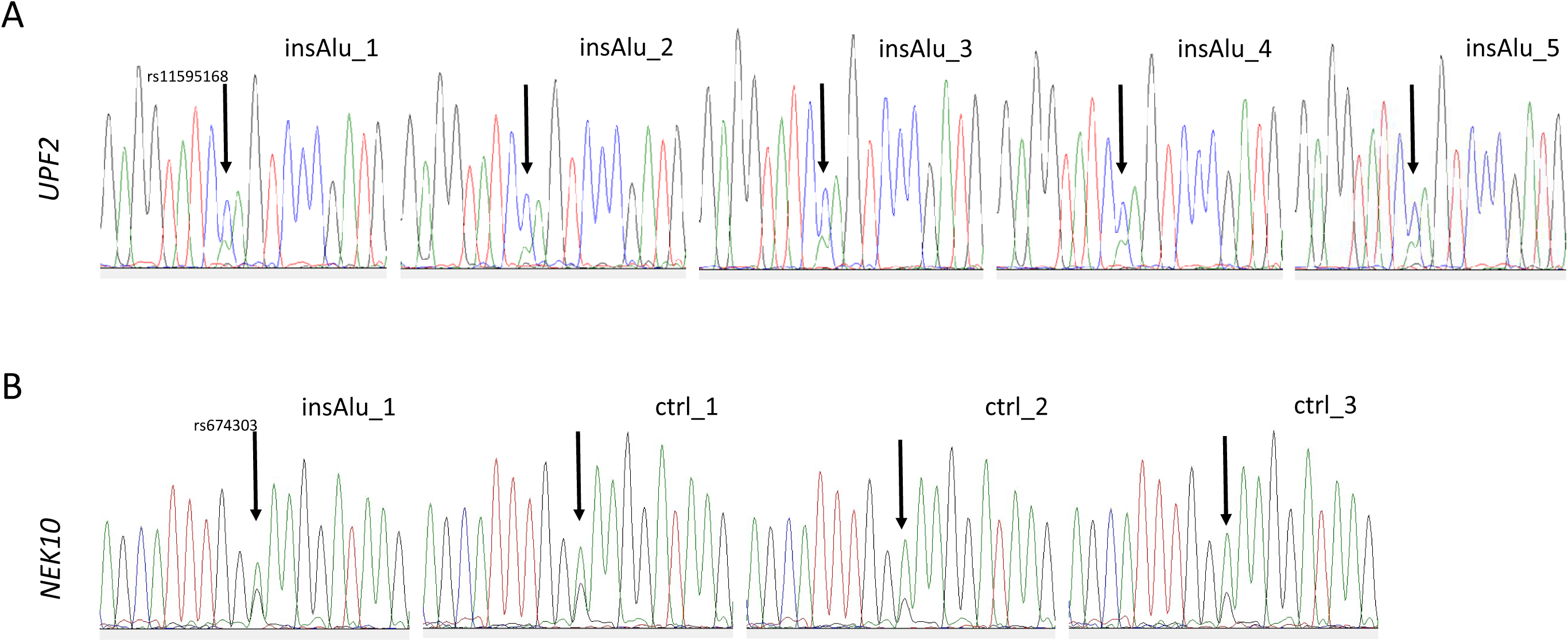
Title: Alu element insertions in *UPF2* and *NEK10* do not induce allelic up or down regulation. Legend: Sanger sequencing experiments were performed on RNA material obtained from blood leucocytes. The chromatograms show a nucleotide region containing a polymorphic position (a SNP, indicated by an arrow) for which all tested samples were heterozygous at the genomic level. (A) In addition to be heterozygous for c.1539A>C (rs11595168), all 5 samples were also heterozygous for the Alu insertion in the intronic region of *UPF2* (c.3408+98_+111insAlu). The c.1539C allele is very rare and was not detected among available control samples. (B) In addition to be heterozygous for c.1674A>G (rs674303), 1 of the 4 samples was also heterozygous for the Alu insertion in the intronic region of *NEK10* (c.26+891_908insAlu).

**Figure 4:**
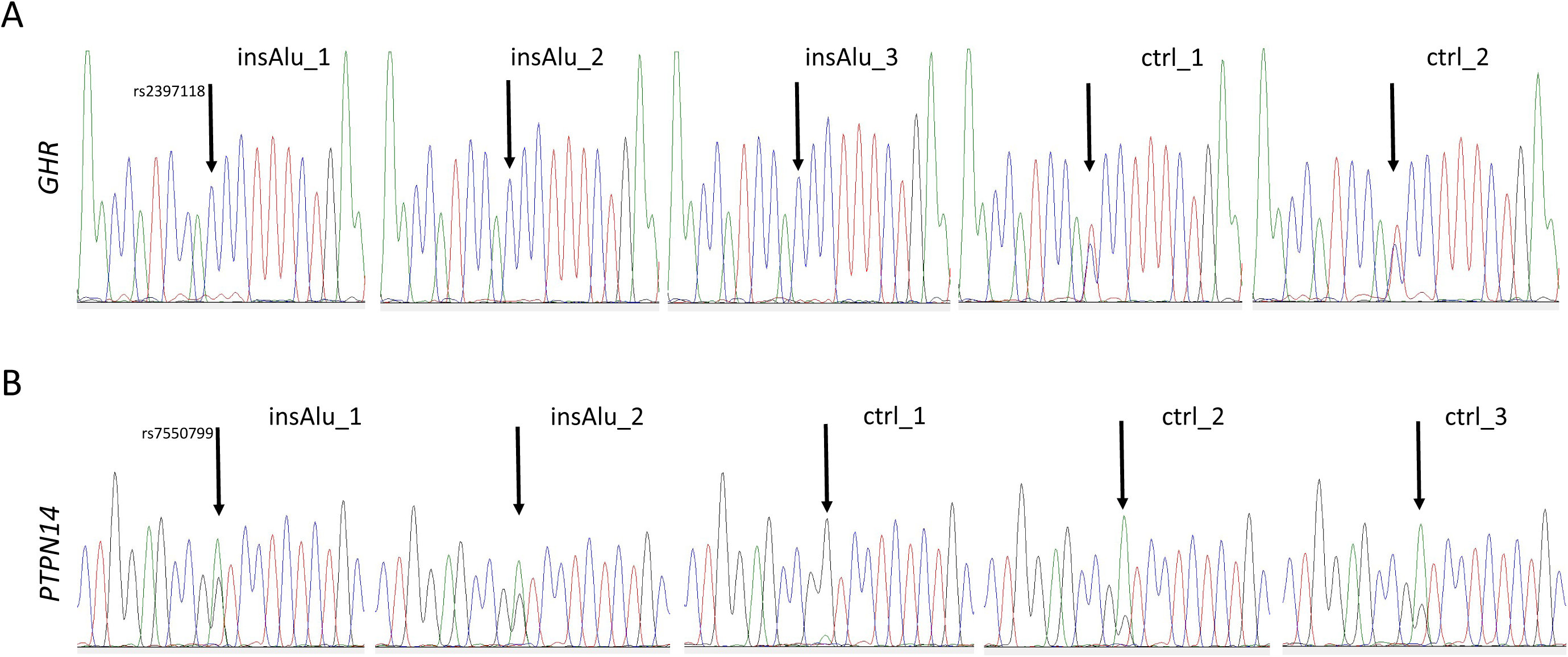
Title: Alu element insertions in *GHR* and *PTPN10* modulate allelic expression. Legend: Sanger sequencing experiments performed on RNA material obtained from blood leucocytes. The chromatograms show a nucleotide region containing a polymorphic position (arrow) for which the tested samples were heterozygous at the genomic level. (A) In addition to be heterozygous for c.558A>G (rs2397118), 3 of the 5 samples were also heterozygous for the Alu insertion in the 3’UTR of *GHR* (c.*+413_+422insAlu). Allele silencing is observed in all Alu mutation carriers. (B) In addition to be heterozygous for c.978A>G (rs7550799), 2 of the 5 samples were also heterozygous for the Alu insertion in the intronic region of *PTPN14* (c.10066+35_+44insAlu). Note that the two Alu mutation carriers express both alleles well, while one control sample (sample *PTPN14*_ctrl1) expresses almost exclusively the G allele, the 2 other control samples showing a deficit in the G allele.

### PolyA containing insertions without identifiable TSD

In addition to the 18 high confident retrotransposition mediated insertions for which we could identify a TSD (described above), our IN SILICO pipeline also identified 14 candidate insertion sites for which a TSD could not be deduced using IGV (see Additional file 1 and Table 3). A summary of each of the targeted gene’s function is presented in Additional file 2 (see also Table 4 for PubMed hits). A lower coverage at sequencing most probably explains the problematic discovery and characterization of these polyA containing insertions. However, for eight of them we found parts of retrotransposon sequences (7x Alu, 1x L1) by analyzing the mates of the discordantly mapped mate pairs (Table 3). In addition, 7 out of these 14 expected retrotransposition events map at the same genomic position as do elements reported in the database for Retrotransposition Insertion Polymorphisms in Humans (dbRIP[26]). Consequently 10 out of the 14 polyA containing insertions can be linked to a retrotransposon, making them still strong candidate retrotransposon insertion sites with potential pathological consequences (Table 3). The high rate of confirmed retrotransposon mediated insertions observed among the candidate polyA containing insertions we identified (28 out of 32) is a good indicator for the high specificity of the followed screening strategy.

**Table 3:** List with the 14 polyA associated insertions for which a TSD could not be defined.

| Targeted gene (region) | Insertion type | Position of insertion in cDNA (RefSeq) | Position of insertion in hg19 | Retrotransposon sequence in mate? | # carriers cl_5 | # carriers cl_3 | reported in dbRIP? |
| --- | --- | --- | --- | --- | --- | --- | --- |
| ANAPC16 (3'UTR) | polyA | c.*+397 (NM_001242546.1) | chr10:73993271 | no | 2 | 2 | yes (Alu) |
| ARHGAP1 (intron) | polyA | c.317+71 (NM_004308) | chr11:46709653 | yes (Alu) | 1 | 1 | no |
| AUH (intron) | polyA | c.599-122 (NM_001698) | chr9:94058481 | yes (L1) | 1 | 4 | yes (L1) |
| CLEC1B (intron) | polyA | c.438+306 (NM_016509) | chr12:10149139 | no | 1 | 2 | no |
| DHRS1 (intron) | polyA | c.374+111 (NM_138452) | chr14:24765604 | yes (Alu) | 1 | 1 | no |
| DYNC2H1 (intron) | polyA | c.6139+43 (NM_001377) | chr11:103048592 | yes (Alu) | 1 | 1 | yes (Alu) |
| ETF1 (3'UTR) | polyA | c.*+2210 (NM_004730) | chr5:137841784 | no | 1 | 1 | no |
| GCNT1 (3'UTR) | polyA | c.*+2545 (NM_001490) | chr9:79121129 | yes (Alu) | 1 | 1 | no |
| GSAP (intron) | polyA | c.1491+495 (NM_017439) | chr7:76980766 | yes (Alu) | 1 | 1 | yes (Alu) |
| MEOX2 (intron) | polyA | c.690+95 (NM_005924) | chr7:15666276 | yes (Alu) | 1 | 1 | yes (Alu) |
| MIER1 (3'UTR) | polyA | c.*+1193 (NM_020948) | chr1:67453349 | no | 1 | 1 | no |
| USP38 (intron) | polyA | c.1210-89 (NM_032557) | chr4:144127097 | yes (Alu) | 1 | 1 | yes (Alu) |
| intergenic | polyA | LINC0251 at 46kb | chr6:114017537 | no | 3 | 4 | no |
| intergenic | polyA | SPIN1 at 6,1kb | chr9:91099726 | no | 1 | 3 | yes (Alu) |

Note that for 8 of the 14 polyA containing insertion sites, a partial retrotransposon sequence could be traced in the mate of at least one discordantly mapped read upon IGV inspection (5<sup>th</sup> column). Screening the database for Retrotransposition Insertion Polymorphisms in Human (dbRIP) at the corresponding positions confirmed the presence of a retrotransposon at 7 of the 14 positions (8<sup>th</sup> column).

**Table 4:** Number of hits obtained in PubMed when searching for the polyA targeted genes.

| PubMed domain | title | title/abstract | title/abstract | title/abstract |
| --- | --- | --- | --- | --- |
| Search terms used | "all gene names" | "all gene names" | "all gene names"<br>AND "cancer" | "all gene names"<br>AND "breast cancer" |
| gene (synonyms)<sup>a</sup> |  |  |  |  |
| ANAPC16 (APC16, C10orf104, CENP-27, FLJ33728, bA570G20.3) | 5 | 5 | 0 | 0 |
| ARHGAP1 (CDC42GAP, ~~RhoGAP~~, p50rhoGAP) | 31 | 93 | 15 | 4 |
| ~~AUH~~ | - | - | - | - |
| CLEC1B (CLEC2) | 9 | 49 | 6 | 0 |
| DHRS1 (FLJ25430, MGC20204, SDR19C1) | 1 | 4 | 1 | 0 |
| DYNC2H1 (DHC1b, DHC2, DNCH2, DYH1B, hdhc11) | 26 | 86 | 4 | 0 |
| ETF1 (~~ERF~~, ERF1, RF1, SUP45L1, TB3-1) | 185 | 845 | 21 | 5 |
| GCNT1 (C2GNT, NACGT2, NAGCT2) | 8 | 124 | 35 | 1 |
| GSAP (LOC54103, ~~PION~~) | 6 | 31 | 1 | 0 |
| MEOX2 (GAX, ~~MOX2~~) | 85 | 254 | 22 | 1 |
| MIER1 (KIAA1610, MI-ER1, hMI-ER1) | 10 | 15 | 2 | 2 |
| USP38 (HP43.8KD, KIAA1891) | 6 | 10 | 3 | 1 |
| intergenic: LINC0251 at 46kb | 0 | 0 | 0 | 0 |
| intergenic: SPIN1 at 6kb (~~SPIN~~, TDRD24) | 18 | 34 | 14 | 4 |

This table shows the number of hits obtained in PubMed when using as search term a specific gene or its synonym (= "all gene names") in combination with "cancer" or "breast cancer" and when restricting the search to particular domains (title or abstract). <sup>a</sup>gene names generating excessive number of false positive hits were removed (strikethrough). Search was performed on 14/01/2020

## DISCUSSION

De novo retrotransposition events occurring within protein coding regions are expected to be very rare as they would strongly affect protein integrity, most often in a detrimental manner [5, 20, 21]. Our results fully match with these earlier observations as the only rare polymorphic retrotransposon that inserted exome wide into a coding sequence among our 65 patients (positive BRCA1 control excluded) was an Alu element into the *ZNF442* gene (a short summary of the function of all genes targeted with a high confident retransposon mediated insertion is presented in Additional file 3). A mutation specific PCR screen (see Table 2) detected this variant in only one out of 710 familial BC patients. Since this patient has North African roots, we cannot exclude that the mutation is more prevalent in that region. *ZNF442* has been poorly investigated till now (see outcomes of PubMed searches for the different targeted genes in Table 5) although the gene has been reported as a good candidate driver gene for the neoplastic process of breast and colorectal cancer[27]. The second identified retrotransposition mediated event that destroys protein integrity is the UQCR10 transcript inserting into the *C1orf194* coding sequence (creating a *UQCR10* processed pseudogene). This polymorphic insertion has been previously observed[7, 28] and was present in 7 of our 65 samples. The mutant allele is unable to produce normal C1orf194 protein, but we cannot exclude that the *UQCR10* pseudogene is expressed as a recombinant mRNA leading to overproduction of UQCR10 protein since a near full length UQCR10 transcript (from c.-15 till polyA tail) inserted in the 5’ coding region (c.58_70) of the *C1orf194* gene in the same orientation (Table 1). Little is known about these two genes, but none has been linked to breast cancer yet. Interestingly, two seemingly dominant heterozygous missense mutations in *C1orf194* were recently associated with Charcot-Marie-Tooth disease[29], the most common form of inherited peripheral neuropathy. The authors reported a Ca^2+^ regulatory function for C1orf194 and therefore suggested that this gene (and consequently any mutated form at it) may also be associated with other neurodegenerative disorders characterized by altered Ca^2+^ homeostasis.

**Table 5:**
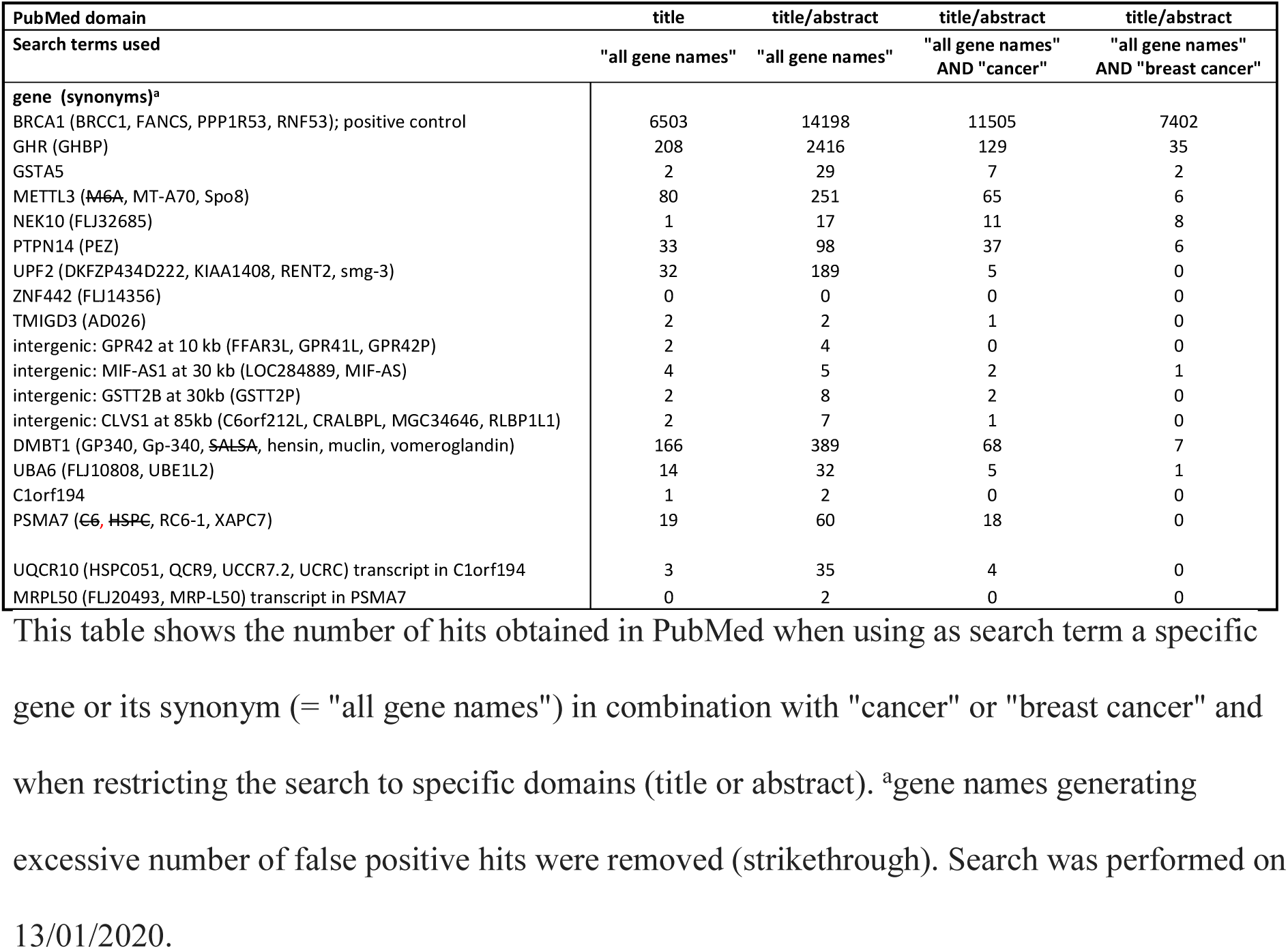
Number of hits obtained in PubMed when searching for high confidence targeted genes.

The large majority of high confident retrotransposon insertions we could identify were located in non-coding regions (15 out of 17). However, such regions are not the primary targets when performing WES and consequently their coverage at sequencing will often be (very) low, or they will not be covered at all. Therefore, we believe much more intronic (near exonic) and UTR located polymorphic retrotransposons escaped our attention, the 15 ones reported herein representing the tip of an iceberg. The majority of the identified retrotransposon insertions incorporated into intronic sequences, close to the intron-exon boundaries. The genes targeted by these retrotransposition events can be linked to several biochemical processes (Additional file 3) that might be involved in breast cancer development: they can have a tyrosine kinase activity (*NEK10*[30, 31]), a Tyrosine phosphatase activity (*PTPN14*[32]), being involved in the ubiquitination process (*UBA6*[33]), in the posttranscriptional methylation of internal adenosine residues in eukaryotic mRNAs (*EMTTL3* [34, 35]), in the regulation of the nonsense mediated decay pathway (*UPF2*[36]), have a suppressive role in osteosarcoma progression (*TMIGD3* [37]) or are considered as candidate tumor suppressor gene for different cancer types (*DMBT1* [38-40]). Interestingly, four of the five intronic Alu insertions occurred in antisense orientation, suggesting that these insertions might contribute to the generation of alternative transcript forms[41]. We also identified two retrotransposon insertions in 3’UTR gene regions (in *GHR* and *HSD17B12*), one in the promoter of *GSTA5* (a glutathione S transferase catalyzing the conjugation of reduced form of glutathione to xenobiotic substances for the purpose of detoxification) and three in intergenic regions (in one case with nearest gene linked to BC: *MIF-AS1*[42]). oMre convincing for their role in breast cancer predisposition is that 3 retrotransposons inserted into a gene that had been previously associated to breast cancer risk (*GHR*[43-45], *DMBT1* [46-48] and *NEK10*[49, 50]). The potential cancer risk associated to these particular mutations should be confirmed by investigating a much higher number of BC cases and controls.

Furthermore, we could successfully investigate the consequences of four different Alu element insertions in non-coding gene regions at the molecular level by determining the relative expression level of the wild type versus mutant allele in leucocytes. Analysis of the Alu insertions in NEK10 and UPF2 did not reveal a marked decrease or increase of the mRNA levels generated from the retrotransposon targeted alleles. Nevertheless, a pathogenic (or protective) contribution by these mutated alleles cannot be excluded yet, as they may for instance generate splicing alterations that cannot be detected with the performed test or may result in cell type specific effects not observed in leucocytes. oMreover, the recently revealed mechanism of onco-exaptation[51, 52] whereby cryptic regulatory elements within transposons can be epigenetically reactivated in precancerous cells may also drive cancer risk. In sharp contrast, the Alu insertion in the 3’UTR of *GHR* clearly resulted in allele silencing. Since several association studies indicated that *GHR* is implicated in breast cancer predisposition[43-45], it is tempting to designate the Alu insertion in *GHR* as (one of) the causal mutation(s) for the reported cancer risk modulation. The interpretation of the data obtained when investigating the Alu insertion in *PTPN14* was more challenging since (near) mono allelic expression was already observed in the three patients that did not carry the germline Alu insertion in PTPN14. Fortunately, literature digging revealed a table wherein PTPN14 was listed as a high-confidence imprinted human gene candidate with maternal expression[24]. The LOI observed in our two Alu carriers suggests that the integration of one single Alu element into the imprinted PTPN14 gene allele would be sufficient to fully reactivate this allele[25]. The observed LOI also suggest that both patients inherited the mutated PTPN14 allele from their father (could not be verified) and that in case the mutated allele is maternally inherited, LOI will not be observed. Since *PTPN14* was recently identified as a mediator of the tumor suppressor activity of *p53* and regularly mutated in cancer[32], its imprinted nature and the loss of imprinting (LOI) induced by particular mutations are two elements that contribute additional levels of complexity to the molecular mechanism leading to cancer when the *PTPN14* gene is involved.

## CONCLUSIONS

Our study indicates that raw WES data obtained from clinical samples (whose availability is exponentially growing) hide a manageable number of retrotransposition mediated polymorphic mutations that can be dug up when using appropriate IN SILICO tools. A significant fraction of these mutations will affect gene function, even when located outside protein coding regions, and consequently may be (one of) the pathogenic factor(s) responsible for a patient’s disease. Our investigation was restricted to 65 non BRCA1/2 familial BC patients, but much more patients should be analyzed (in future genome wide) to obtain a broader view of all possible retrotransposon mediated insertions involved in the disease, and to determine their respective molecular and pathogenic consequences. oMreover, transposon mediated insertions resulting in modified breast cancer risk may also generate other clinical phenotypes. For example, the growth hormone receptor (which is encoded by the *GHR* gene) is expressed in a broad range of tissues and involved in fundamental biological processes such as growth promotion, metabolism, cell division and regeneration[53], the most typical clinical syndrome associated to GHR deficiency being dwarfism (Laron syndrome). Accordingly, the clinical consequences associated to the presence of an Alu insertion in the 3’UTR of *GHR* (which leads to allele silencing) are most probably not limited to breast cancer risk. As pathogenic retrotransposon insertions are not limited to the familial BC syndrome, the described mutation screen (or an alternative version of it, e.g. for the analysis of WES data obtained with the exome capture kit from Agilent) should be applied for all diseases with a suspected genetic etiology. Indeed, screening patients with a specific genetic disease will enrich for insertions in genes involved in that disease (in preparation). Centralized databases registering all identified polymorphic retrotransposition events should be further expanded[7, 26, 54], with inclusion of population specific allele frequencies, as is the case for classical variations. In order to identify good candidate pathogenic Alu element insertions, Payer et al.[55] used the outputs of GWAS studies to restrict their search to genomic loci previously implicated in disease risk. Conversely, our data indicate that microchips should be developed allowing genome wide genotyping for the presence of this particular type of polymorphic insertions, in order to perform retrotransposition specific GWAS’s for a multitude of diseases or traits. Finally, the genome wide identification and investigation of polymorphic retrotransposon insertions in clinical samples will not only lead to a better understanding of diseases, but will also contribute to elucidate more basic genetic mechanisms such as gene imprinting, and will help to evaluate the impact that de novo retrotransposition mediated insertions still have on human genome evolution.

## METHODS

### Patient Material

A WES study using the SeqCap EZ Exome v3.0 kit from Roche has previously been performed using blood samples from probands of 65 unrelated high risk BC families with the aim to identify classical cancer predisposing mutations (protein truncating SNPs and Indels, and splice site mutations)[17]. All patients were recruited at the UZ-Brussel hospital and met the requirements for a diagnostic *BRCA1* and *BRCA2* mutation analysis. All except one (the positive control: c.1739_1740insAlu in *BRCA1*[13]) were negative for a pathogenic *BRCA1* or *BRCA2* mutation. A large subset of these 65 patients (57) belongs to families with at least two first degree blood relatives with BC. The FastQ files generated during this previous study were reused for the present study. The genomic DNA from the positive control was resubmitted for WES analysis using the SureSelect Human All Exon V6 kit from Agilent (performed by BGI Genomics Co, China) to compare both WES approaches. In addition, blood samples obtained from 710 probands from independent non BRCA1/2 BC families (including the 65 probands used for the WES analysis) were used for a PCR based genomic screen to identify additional carriers of Alu element insertions in case insufficient samples were available for RNA analysis.

### Strategy for retrotransposon detection using WES-BAM files

Binary alignment files (BAMfiles) are outputs (binary versions) of a short read aligner (e.g. BWA[56]) that can map the read pairs generated during the sequencing process (paired-end sequencing) towards a reference sequence, for instance the reference human genome (hg19). In particular circumstances, these read pairs are discordantly mapped while generated from the same small genomic fragment, meaning that the mate reads map at very different positions on the reference genome.

To trace retrotransposon insertions not present in the human reference sequence we used, in a first try-out, the open-source RetroSeq software described by Keane et al.[18] as this software can be run on virtually any computer[57]. This software needs as input a BAMfile (we used the WES BAMfile generated with a positive control sample harboring the c.1739_1740insAlu mutation in *BRCA1*; the library was prepared using the exome capture kit SeqCap EZ Exome v3.0 from Roche), a reference genome (hg19) and a library of mobile element sequences. RetroSeq operates in two phases, the first being the discovery phase where discordantly mapped (and one end mapped) mate pairs are screened for the presence of a mobile element and in the affirmative assigned to a particular type of mobile element. In the second phase (calling phase), the sequence of the anchoring mate read is used to localize the polymorphic mobile element on the reference genome. Unfortunately, when applying RetroSeq on our positive control sample we could not detect the Alu insertion in *BRCA1*. RetroSeq is primarily designed to identify non reference mobile elements in Whole Genome Sequencing data, while companies that design kits for exome enrichment try to avoid the capture of nucleotide fragment harboring repetitive sequences. To circumvent this obstacle, we decide in a first step to search not for mobile elements but for polyA stretches (settings: minimum 80 nucleotides long; 90% ID) within the discordantly mapped reads of each patient data set using the first module of the RetroSeq software (discovery phase). Among the discordantly mapped read pairs traceable in BAMfiles we expect to find those generated from genomic DNA fragments that contain both non reference retrotransposon sequences and exon (flanking) sequences (see Figure 1). By selecting discordantly mapped reads containing a long polyA stretch in one of the paired reads, we will only retain the junctions at the 3’ end of the retrotransposon. The location of the mate of each polyA containing read identified during the discovery phase is recorded in a BED file by the RetroSeq program (together with additional information). This file can be recovered as a .tab file for further analysis (see Additional file 4 for the .tab file generated with the positive control sample saved in Excel).

If coverage is sufficient, several anchor reads (the mates of the polyA containing reads) are expected to align in the same genomic region. Clusters of such reads are subsequently generated when they occur within a 300 bp long genomic interval and a maximum inter read distance of 200 bp. In a next step, each cluster is represented by the anchor read expected to be closest to the integration site (TSD in case of retrotransposons) while the number of reads within each cluster is recorded in a separate column (duplicate reads are removed). To perform these cluster calculation steps, the patient specific .tab file generated by RetroSeq is saved in Excel format, non-relevant information is first deleted and the remaining data are pasted in the first sheet of a preformatted Excel file (provided as Additional file 5). The detailed procedure for cluster calculation is described in next section. We considered clusters with three or more anchor reads as potential candidate insertion sites and retained them for further investigation. When applying this detection strategy to our 65 BC patients, we obtained on average 987 such clusters for each individual. The positive control sample (see Additional file 6) shows 641 clusters with three or more units, with a clear cluster of 17 units at the level of the Alu insertion site in *BRCA1* (TSD is at chr17:41245809-41245825).

To identify and in a second step discard the clusters (genomic regions) picked up in a large fraction of individuals (e.g. false positives resulting from technical artefacts, or highly recurrent polymorphic insertions not present in the reference genome), the Excel outputs obtained from each BC patient were pooled (keeping patient ID tracked) and clusters were again generated. Candidate genomic insertion sites detected in more than 10% of the samples (corresponding to clusters with more than 6 units for the BC cohort) were identified and subtracted from the patient specific output data. After this filtering step, on average 9 clusters with 5 or more reads are obtained for each individual. Visual inspection using the IGV software revealed that the majority of the remaining candidate insertion sites are generated by the presence of a polyA stretch in the reference sequence that resulted in DNA polymerase slippage during the NGS process. Consequently, these candidate genomic insertion sites were manually tracked and listed, and used for subsequent filtering. Additional file 7 provides a preformatted Excel file to perform the combined filtering steps. After filtering, only half of the BC patient samples presented a candidate insertion event (31/65). One patient presented 4 such events (Additional file 1). To minimalize the possibility that these candidate insertion sites would be missed in a subset of individuals because of poorer coverage, all genomic positions identified as insertional target sites during the first screening round in any individual (positions deduced from the presence of a cluster with minimum 5 anchor reads) were re-screened for the presence of a smaller cluster (3 or 4 reads) in all individuals (Additional file 1, columns M&N). Further IGV inspection allowed the identification of a TSD in a significant fraction (18/32) of the obtained candidate insertion sites, strongly suggesting that a retrotransposition event had occurred at those genomic locations (Additional file 1, column K). For validation purposes, the presence or absence of each of these 18 insertions was verified in each of the 65 patients using IGV (Table 1). The recovery rates obtained IN SILICO and manually using IGV matched very well except for two insertions (the *SET* transcript in *DPP10* and an Alu element in *HSD17B12*), the discrepancies being explainable by the poor coverage at sequencing. For the identification of the inserted sequences, we used the Dfam database of repetitive DNA families[58] (https://dfam.org/home) and the Basic Local Alignment Search Tool (https://blast.ncbi.nlm.nih.gov/Blast.cgi) (Additional file 1, column I). As only small stretches of nucleotides (about 50 bp long) from the 5’ end of the inserted sequences can be deduced from the WES generated data, it was not possible to further define the sub-family of the identified retrotransposons.

A screenshot showing read visualization by IGV of the region were the Alu insertion occurred in the positive control sample (*BRCA1*) when using the SeqCap EZ Exome v3.0 kit from Roche for exome capture is presented in Figure 2. A corresponding screenshot showing read visualization when using the SureSelect Human All Exon V6 kit from Agilent is presented in Figure 5. Note that very few discordantly mapped reads are observed when using the Agilent kit, indicating that the detection strategy we described here above is unable to detect retrotransposon insertions when using the WES BAMfiles generated with this kit, although several other identifiers of an Alu insertion are clearly present and useable for the development of an adapted IN SILICO pipeline.

**Figure 5:**
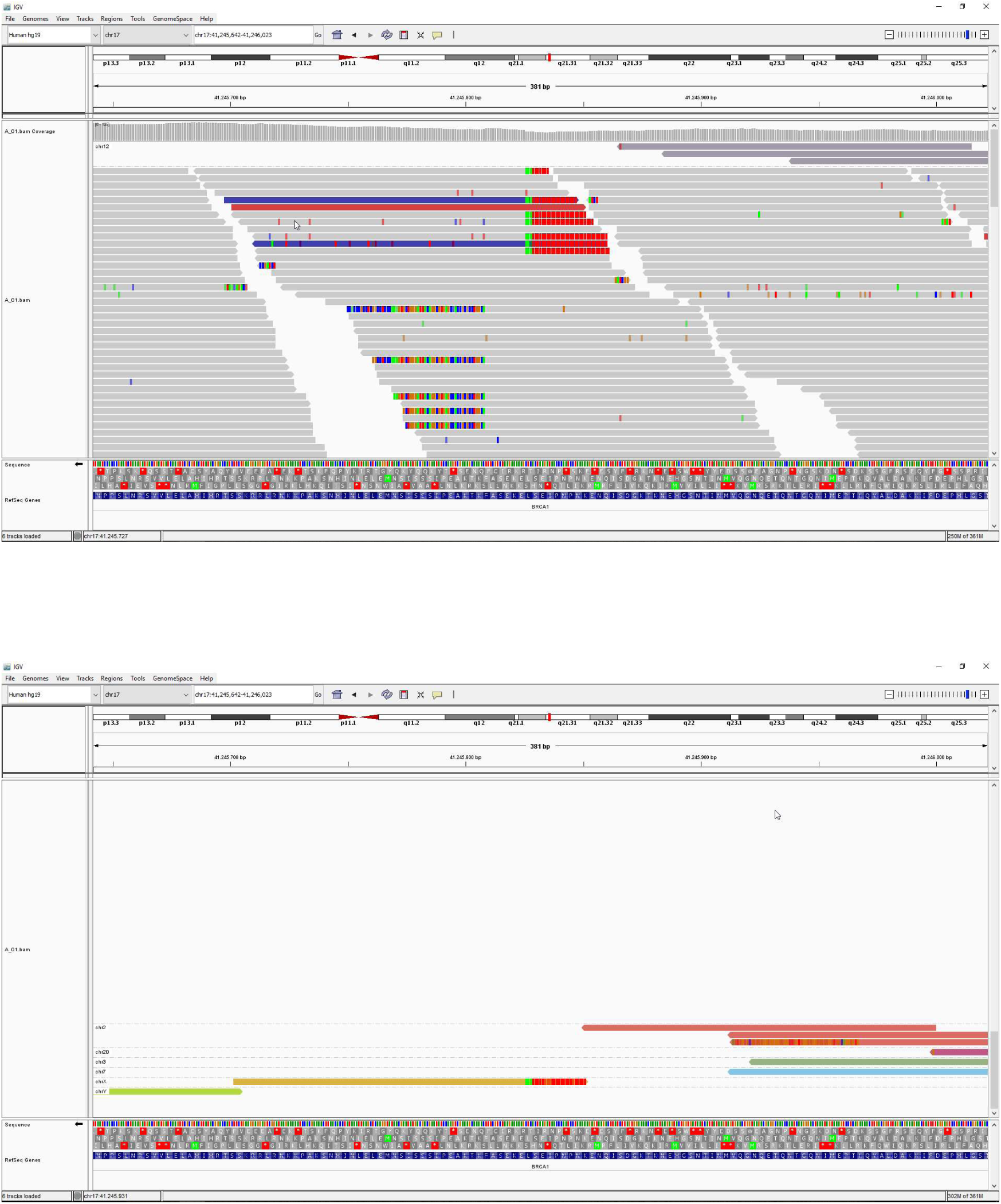
Title: IGV visualization of the control Alu insertion using BAMfile generated with the Agilent kit. Legend: Screenshots of IGV outputs restricted to the genomic region where the Alu element insertion occurred (c.1739_1740insAlu in *BRCA1*) in the positive control sample. The BAMfile used for IGV visualization was generated from WES data obtained with the SureSelect Human All Exon V6 kit (exome capture kit) from Agilent. Group alignment is by chromosome of mate. Note that this exome capture kit generates much less discordantly mapped read pairs than the kit from Roche (2 versus 22, compare with Figure 2). Accordingly, the required number (five) of discordantly mapped read pairs containing a long polyA track will not be reached and the insertion will not be revealed with our detection strategy. Note that all other characteristics observed for a retrotransposon insertion when using the exome capture kit from Roche (see Figure 2) are also observed when using the kit from Agilent.

### Procedure for cluster calculation starting with the data obtained from RetroSeq

Save the RetroSeq generated tab. file in Excel format and delete the 2 last columns (the .tab file generated with the positive control sample is provided as “Additional file 4”).

Remove all lines not referring to an autosome or to chX (all samples are from females). Sort (A>Z) according to sign in last (6^th^) column.

Select/copy all filled cells and paste in Sheet 1 (“calculate cluster”) of the preformatted Excel file for cluster calculation (Additional file 5).

In this Sheet 1, select for “TRUE” in column I (OR) and select/copy all filled cells in columns A to J.

Paste the data in Sheet 2 (“simplify cluster”), and in column A, order by “color”. Delete the entire rows corresponding to the cells with pink background (this will remove duplicates).

In column O (chr), deselect “blank” and select/copy all informative cells in columns O to U.

Paste these data in Sheet 3 (“all clusters”), and deselect “2” in column G (to keep clusters with minimum 3 reads).

Select/copy all filled cells and paste in Sheet 4 (annotation list, use the first column for sample ID). The positive control sample will generate 641 clusters (see Additional file 6).

### Procedure to filter out the clusters of secondary interest (false positives or clusters highly recurrent among patients)

Based on the data we obtained from our 65 samples, we generated a table (first sheet of Additional file 7) listing (a) all clusters with minimal 3 reads that were observed in 6 or more of our patients (= clusters present ∼10% or more of the samples) and (b) all clusters that resulted from the presence of a polyA stretch in the reference sequence. This table was subsequently used to identify and discard the less relevant clusters from the patient specific cluster lists (as described above). This list is provided with the only intension to familiarize with the described procedure. The content of this list can depend on the type and number of samples used for its generation (type of disease, ethnicity…) but also on the followed wet and dry bench procedures.

Select/copy all filled cells of Sheet 4 (annotation list) obtained from a patient specific cluster calculation (see above).

Open Sheet 1 (comm65>=6 + sample) of the preformatted Excel file for cluster filtering (Additional file 7) and paste the data (in column A) down the preexisting list.

Apply Sort (A>Z) on column C (start) and thereafter on column B (chr).

Select/copy the generated dataset and paste in Sheet 2 (calculate cluster_com65>=6).

Select for “0” in column L (“cluster”), for “FALSE” in column K (“OR”) and for “sampleID” in column A.

Select/copy/paste clusters with five or more reads to the “results” Sheet 3.

The obtained candidate insertion sites (represented by each of the generated clusters in Sheet 3) can be manually validated by IGV inspection of the corresponding BAMfiles at the indicated positions.

### Hemi-nested PCR’s for genomic validation and screen

All uncommon Alu element insertions identified with high confidence (presence of a TSD) within or close to exonic gene sequences or promoter sequences (7 in total, see Table 1) were validated by a 2 step PCR. During the first step, and starting with an input of 6.25 ng genomic DNA, primers flanking the suspected integration site are used to generate a 200-300 bp long PCR fragment. This fragment is used as template (after 2000x dilution) for the second step of the hemi-nested PCR wherein one primer of the first step PCR is used in combination with an Alu specific primer. The first step of the nested PCR should work for all genomic DNA samples. The second step will work only when the patient is carrier of the targeted Alu insertion and the appropriate primer combination is used. Both PCR steps were run on a real-time PCR instrument (LightCycler 480 II from Roche). DNA samples from Alu carriers (and negative controls) according to our WES screen were used for validation. During the first step, an amplification with reproducible Ct and melting curve was observed with all samples while during the second step only samples with an Alu insertion gave an amplification signal (with reproducible Ct and melting curve). The obtained PCR fragments were further evaluated by agarose gel electrophoresis and all showed the expected size. The primers used for PCR amplification are shown in Table 6. The Alu specific primer (Alu/Rev) is located in a well conserved region and points to the 5’ extremity of the transposon. Consequently, this primer will never allow amplification of the 3’ extremity (polyA tail) of the transposon. The choice of the second primer for the second step of the PCR will depend on the orientation of the Alu element compared to the orientation of the targeted gene (see Table1, 2^nd^ column). In order to identify additional carriers of the Alu insertions characterized during the raw WES data screen, the same nested PCRs were performed on genomic DNA from 710 familial BC patients (Table 2).

**Table 6:**
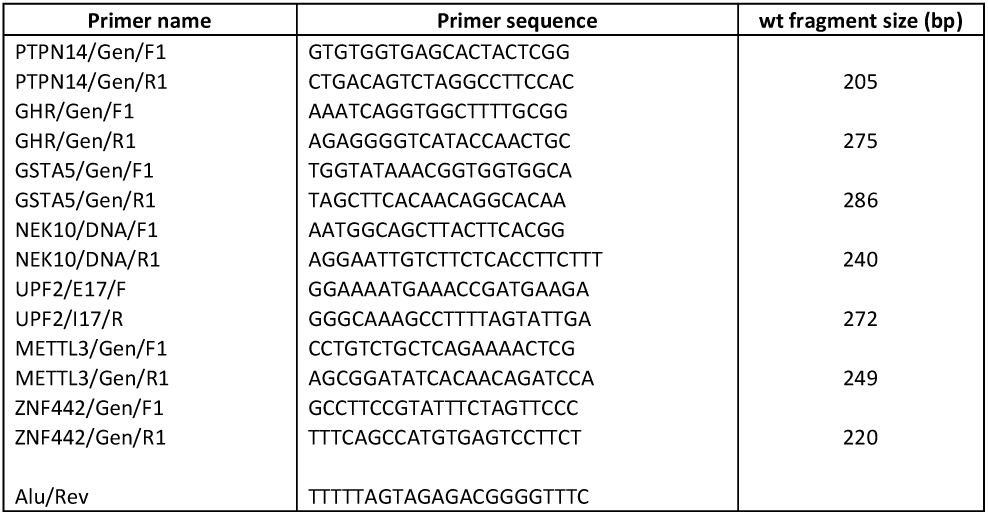
Primers used for the detection of 7 Alu element insertions in genomic DNA

### Genotyping at polymorphic sites

To investigate whether an Alu element that integrated into the non-coding regions of a gene affects expression levels of the targeted allele, a polymorphic site allowing allelic discrimination at the cDNA level must first be identified. This was done by IGV inspection of the BAMfiles from patients that had revealed a polymorphic Alu insertion during our screen (Additional file 1). To increase the number of samples that could be enrolled in the allele expression study, the patients whose genomic DNA revealed an interesting Alu insertion upon mutation specific PCR analysis were genotyped for the corresponding polymorphic site. PCR primers are presented in Table 7. Heterozygosity was determined by High Resolution eMlting Curve Analysis.

**Table 7:**
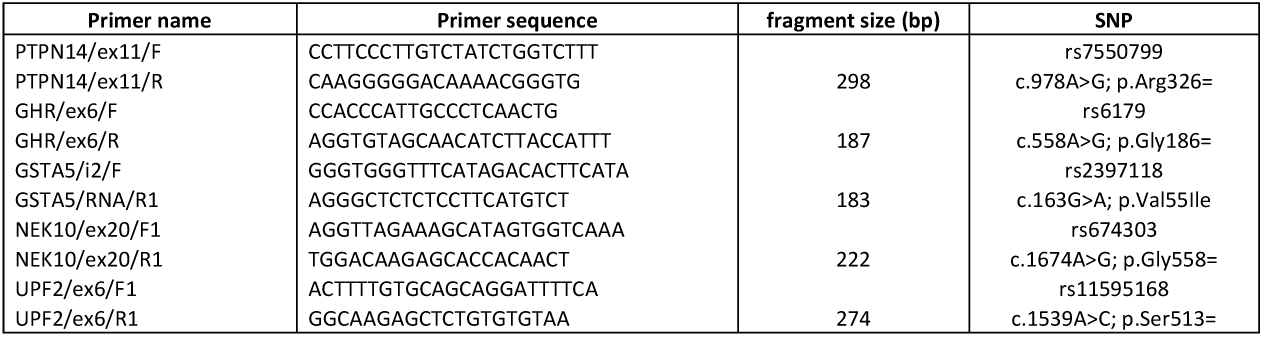
Primers used for genotyping at polymorphic positions (for RNA study)

### RNA isolation, cDNA synthesis and RT-PCR

RNA isolation was performed using the RNeasy iMni kit (Qiagen) with additional DNAse step using manufacturer’s protocol. cDNAs were prepared using Superscript II reverse transcriptase (Life Sciences) using hexamer random primers and qPCR was performed using the SybrGreen aMster kit (Roche) on a LightCycler 480 II. Primers are presented in Table 8.

**Table 8:**
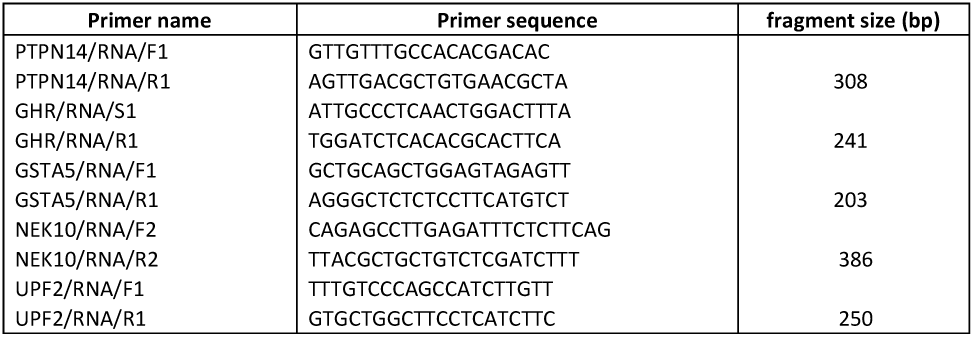
Primers used for cDNA amplification (and Sanger sequencing)

### Sequencing

Purified PCR fragments were Sanger sequenced by the VIB Genetic Service Facility, from the University of Antwerp.

## DECLARATIONS

Ethics approval and consent to participate: Patient recruitment and blood sampling were performed according to the ethical procedures approved by the institutional ethics committee of the UZ Brussel. For the concerned cases, peripheral blood was collected after obtaining a written informed consent for a broad genomic analysis covering also incidental findings in genes predictive for other diseases.

### Consent for publication

Not applicable

### Availability of data and materials

a dataset allowing the reproduction of the outputs for a positive control is provided as “additional file”. Additional datasets are available on reasonable request

### Competing interests

the authors declare that they have no competing interests

### Funding

financial support from “Kom Op Tegen Kanker” is acknowledged

### Authors’ contribution

SDB and ET conceived the project, designed and performed the experiments, interpreted the data and wrote the manuscript. JDG is responsible for patient recruitment and counselling. All authors reviewed and approved the manuscript.

#### Acknowledgements

not applicable

### Authors’ information (optional)

not applicable

to be added to ref35 (bold): r…ole of RNA m **6A Methylation in Cancer**. (not accepted by EndNote)

**FIGURES (titles and legends)**

Strong candidate integration sites are retained when generated by clusters counting at least 5 polyA containing discordantly mapped read pairs, when present in less than 10% of the samples, and when a TSD is deducible upon IGV inspection in at least one patient. ^a^ indicates the number of nucleotides available to identify retrotransposon type (using the Dfam database [58]). As only about 50 nucleotides are available for transposon identification, it was not possible to define sub-classes. ^b^ indicates the number of patients (out of 65) who carry the mutation based on clusters counting at least 5 discordantly mapped read pairs. ^c^ indicates the number of patients who carry the same mutation based on clusters counting at least 3 discordantly mapped read pairs. ^d^ indicates the number of patients harboring the same mutation based upon IGV inspection. Note that for two insertions (SET in *DPP10* and Alu in *HSD17B12*) a much higher number of carriers is revealed upon IGV inspection. This discrepancy results from the poor coverage at sequencing. ^e^ indicates the Alu insertions for which a genomic hemi-nested PCR has been developed and applied for validation (see Methods).

## ADDITIONAL MATERIAL > INFORMATION LIST

Additional file 1 (Excel file)

Title: Strong candidates retrotransposon mediated insertion sites

Description: This table lists all strong candidate insertion sites detected with the RetroSeq software (minimum 5 discordantly mapped read pairs pointing to the same genomic location) and further filtered using Excel software to remove false positives and insertions occurring in more than 10% of the investigated patient samples.

Additional file 2 (Word file)

Title: Retrotransposon targeted genes with no TSD identified

Description: For each gene presumably targeted by a retrotransposition event that could not be confirmed by the presence of a TSD, a short description is given. The gene/protein descriptions provided by “Entrez Gene Summary” (https://www.ncbi.nlm.nih.gov/gene), by “GeneCards Summary” (https://www.genecards.org) and by “UniProtKB/Swiss-Prot Summary” (https://uniprot.org) are presented, when available.

Additional file 3 (Word file)

Title: Retrotransposon targeted genes with TSD identified

Description: For each gene targeted by a retrotransposition event that could be confirmed by the presence of a TSD, a short description is given. The gene/protein descriptions provided by “Entrez Gene Summary” (https://www.ncbi.nlm.nih.gov/gene), by “GeneCards Summary” (https://www.genecards.org) and by “UniProtKB/Swiss-Prot Summary” (https://uniprot.org) are presented, when available.

Additional file 4 (Excel file)

Title: RetroSeq generated list of anchor reads when using the positive control sample

Description: This table lists all discordantly mapped read pairs (19.674 in total) selected by the RetroSeq software when analyzing the positive control sample (patient harboring the c.1739_1740insAlu mutation in BRCA1). The input BAMfile was generated using the exome capture kit SeqCap EZ Exome v3.0 from Roche for library preparation. To be selected, one read of the discordantly mapped read pair must contain a long polyA stretch, while the position of the mate of this read on the reference genome is used for ordering the RefSeq selected read pairs.

Additional file 5 (Excel file)

Title: Excel file for cluster calculation

Description: This pre-formated Excel file can be used to extract potential retrotransposon insertion sites from lists generated with the RetroSeq software (e.g. the list presented in Additional file 4)

Additional file 6 (Excel file)

Title: Output of cluster calculation for the positive control sample

Description: This table lists all potential retrotransposon integration sites observed in the positive control sample (641 in total). Integration sites are retained only when 3 or more anchor reads cluster in the same genomic interval (max 300bp long).

Additional file 7 (Excel file)

Title: Excel file for cluster filtering

Description: This pre-formated Excel file allows further filtering of the potential retrotransposon integration sites (output of Additional file 5). Highly recurrent integration sites (occurring in more than 10% of the investigated population) or false positives are removed during this filtering step.

## Notes

### Competing Interest Statement

The authors have declared no competing interest.

