## Additional file 2 for "A systematic screen of breast cancer patients’ exomes for retrotransposon insertions reveals disease associated genes"

### **Additional file 2:** A short description as provided by “Entrez Gene Summary” (https://www.ncbi.nlm.nih.gov/gene), by “GeneCards Summary” (https://www.genecards.org) and by “UniProtKB/Swiss-Prot Summary” (https://uniprot.org) is given, if available, for each gene presumably targeted by a retrotransposition event that could not be confirmed by the presence of a TSD.

**ANAPC16:**

**GeneCards Summary for ANAPC16 Gene**

ANAPC16 (Anaphase Promoting Complex Subunit 16) is a Protein Coding gene. Among its related pathways are Cellular Senescence (REACTOME) and CDK-mediated phosphorylation and removal of Cdc6.

**UniProtKB/Swiss-Prot Summary for ANAPC16 Gene**

Component of the anaphase promoting complex/cyclosome (APC/C), a cell cycle-regulated E3 ubiquitin ligase that controls progression through mitosis and the G1 phase of the cell cycle. The APC/C complex acts by mediating ubiquitination and subsequent degradation of target proteins: it mainly mediates the formation of 'Lys-11'-linked polyubiquitin chains and, to a lower extent, the formation of 'Lys-48'- and 'Lys-63'-linked polyubiquitin chains

**ARHGAP1:**

**Entrez Gene Summary for ARHGAP1 Gene**

This gene encodes a member of a large family of proteins that activate Rho-type guanosine triphosphate (GTP) metabolizing enzymes. The encoded protein contains a SRC homology 3 domain and interacts with Bcl-2-associated protein family members. [provided by RefSeq, Aug 2012]

**GeneCards Summary for ARHGAP1 Gene**

ARHGAP1 (Rho GTPase Activating Protein 1) is a Protein Coding gene. Diseases associated with ARHGAP1 include Exudative Vitreoretinopathy 1 and Exudative Vitreoretinopathy. Among its related pathways are Actin Nucleation by ARP-WASP Complex and Regulation of RAC1 activity. Gene Ontology (GO) annotations related to this gene include GTPase activator activity and SH3/SH2 adaptor activity. An important paralog of this gene is PRR5-ARHGAP8.

**UniProtKB/Swiss-Prot Summary for ARHGAP1 Gene**

GTPase activator for the Rho, Rac and Cdc42 proteins, converting them to the putatively inactive GDP-bound state. Cdc42 seems to be the preferred substrate.

**AUH:**

**Entrez Gene Summary for AUH Gene**

This gene encodes bifunctional mitochondrial protein that has both RNA-binding and hydratase activities. The encoded protein is a methylglutaconyl-CoA hydratase that catalyzes the hydration of 3-methylglutaconyl-CoA to 3-hydroxy-3-methyl-glutaryl-CoA, a critical step in the leucine degradation pathway. This protein also binds AU-rich elements (AREs) found in the 3' UTRs of rapidly decaying mRNAs including c-fos, c-myc and granulocyte/ macrophage colony stimulating factor. ARE elements are involved in directing RNA to rapid degradation and deadenylation. This protein is localized to the mitochondrial matrix and the inner mitochondrial membrane and may be involved in mitochondrial protein synthesis. Mutations in this gene are the cause of 3-methylglutaconic aciduria, type I. Alternative splicing results in multiple transcript variants. [provided by RefSeq, Sep 2015]

**GeneCards Summary for AUH Gene**

AUH (AU RNA Binding Methylglutaconyl-CoA Hydratase) is a Protein Coding gene. Diseases associated with AUH include 3-Methylglutaconic Aciduria, Type I and 3-Methylglutaconic Aciduria. Among its related pathways are Valine, leucine and isoleucine degradation and Amino Acid metabolism. Gene Ontology (GO) annotations related to this gene include RNA binding and enoyl-CoA hydratase activity. An important paralog of this gene is ECHDC2.

**UniProtKB/Swiss-Prot Summary for AUH Gene**

Catalyzes the conversion of 3-methylglutaconyl-CoA to 3-hydroxy-3-methylglutaryl-CoA (PubMed:11738050, PubMed:12434311, PubMed:12655555). Also has itaconyl-CoA hydratase activity by converting itaconyl-CoA into citramalyl-CoA in the C5-dicarboxylate catabolism pathway (PubMed:29056341). The C5-dicarboxylate catabolism pathway is required to detoxify itaconate, a vitamin B12-poisoning metabolite (PubMed:29056341). Has very low enoyl-CoA hydratase activity (PubMed:7892223). Was originally identified as RNA-binding protein that binds in vitro to clustered 5'-AUUUA-3' motifs (PubMed:7892223)

**CLEC1B:**

**Entrez Gene Summary for CLEC1B Gene**

Natural killer (NK) cells express multiple calcium-dependent (C-type) lectin-like receptors, such as CD94 (KLRD1; MIM 602894) and NKG2D (KLRC4; MIM 602893), that interact with major histocompatibility complex class I molecules and either inhibit or activate cytotoxicity and cytokine secretion. CLEC2 is a C-type lectin-like receptor expressed in myeloid cells and NK cells (Colonna et al., 2000 [PubMed 10671229]).[supplied by OMIM, Jan 2011]

**CeneCards Summary for CLEC1B Gene**

CLEC1B (C-Type Lectin Domain Family 1 Member B) is a Protein Coding gene. Diseases associated with CLEC1B include Human Immunodeficiency Virus Type 1 and Bleeding Disorder, Platelet-Type, 11. Among its related pathways are G beta-gamma signaling through PI3Kgamma and Response to elevated platelet cytosolic Ca2+. Gene Ontology (GO) annotations related to this gene include transmembrane signaling receptor activity and carbohydrate binding. An important paralog of this gene is CLEC12B.

**UniProtKB/Swiss-Prot Summary for CLEC1B Gene**

C-type lectin-like receptor that functions as a platelet receptor for the lymphatic endothelial marker, PDPN (PubMed:18215137). After ligand activation, signals via sequential activation of SRC and SYK tyrosine kinases leading to activation of PLCG2 (PubMed:18955485).

(Microbial infection) Acts as a receptor for the platelet-aggregating snake venom protein rhodocytin. Rhodocytin binding leads to tyrosine phosphorylation and this promotes the binding of spleen tyrosine kinase (SYK) and initiation of downstream tyrosine phosphorylation events and activation of PLCG2 (PubMed:16174766, PubMed:18955485).

(Microbial infection) Acts as an attachment factor for Human immunodeficiency virus type 1 (HIV-1) and facilitates its capture by platelets (PubMed:16940507).

**DHRS1:**

**Entrez Gene Summary for DHRS1 Gene**

This gene encodes a member of the short-chain dehydrogenases/reductases (SDR) family. The encoded enzyme contains a conserved catalytic domain and likely functions as an oxidoreductase. Multiple alternatively spliced variants, encoding the same protein, have been identified. [provided by RefSeq, Nov 2008]

**GeneCards Summary for DHRS1 Gene**

DHRS1 (Dehydrogenase/Reductase 1) is a Protein Coding gene. Diseases associated with DHRS1 include Specific Language Impairment. Gene Ontology (GO) annotations related to this gene include oxidoreductase activity. An important paralog of this gene is HSDL2.

**DYNC2H1:**

**Entrez Gene Summary for DYNC2H1 Gene**

This gene encodes a large cytoplasmic dynein protein that is involved in retrograde transport in the cilium and has a role in intraflagellar transport, a process required for ciliary/flagellar assembly. Mutations in this gene cause a heterogeneous spectrum of conditions related to altered primary cilium function and often involve polydactyly, abnormal skeletogenesis, and polycystic kidneys. Alternative splicing results in multiple transcript variants encoding distinct proteins. [provided by RefSeq, Jan 2010]

**GeneCards Summary for DYNC2H1 Gene**

DYNC2H1 (Dynein Cytoplasmic 2 Heavy Chain 1) is a Protein Coding gene. Diseases associated with DYNC2H1 include Short-Rib Thoracic Dysplasia 3 With Or Without Polydactyly and Short Rib-Polydactyly Syndrome, Majewski Type. Among its related pathways are Phagosome and Intraflagellar transport. Gene Ontology (GO) annotations related to this gene include ATPase activity and motor activity. An important paralog of this gene is DYNC1H1.

**UniProtKB/Swiss-Prot Summary for DYNC2H1 Gene**

May function as a motor for intraflagellar retrograde transport. Functions in cilia biogenesis. May play a role in transport between endoplasmic reticulum and Golgi or organization of the Golgi in cells (By similarity).

**ETF1:**

**Entrez Gene Summary for ETF1 Gene**

This gene encodes a class-1 polypeptide chain release factor. The encoded protein plays an essential role in directing termination of mRNA translation from the termination codons UAA, UAG and UGA. This protein is a component of the SURF complex which promotes degradation of prematurely terminated mRNAs via the mechanism of nonsense-mediated mRNA decay (NMD). Alternate splicing results in multiple transcript variants. Pseudogenes of this gene are found on chromosomes 6, 7, and X. [provided by RefSeq, Aug 2013]

**GeneCards Summary for ETF1 Gene**

ETF1 (Eukaryotic Translation Termination Factor 1) is a Protein Coding gene. Among its related pathways are Viral mRNA Translation and Gene Expression. Gene Ontology (GO) annotations related to this gene include ribosome binding.

**UniProtKB/Swiss-Prot Summary for ETF1 Gene**

Directs the termination of nascent peptide synthesis (translation) in response to the termination codons UAA, UAG and UGA (PubMed:7990965, PubMed:24486019). Component of the transient SURF complex which recruits UPF1 to stalled ribosomes in the context of nonsense-mediated decay (NMD) of mRNAs containing premature stop codons.

**GCNT1:**

**Entrez Gene Summary for GCNT1 Gene**

 This gene is a member of the beta-1,6-N-acetylglucosaminyltransferase gene family. It is essential to the formation of Gal beta 1-3(GlcNAc beta 1-6)GalNAc structures and the core 2 O-glycan branch. The gene coding this enzyme was originally mapped to 9q21, but was later localized to 9q13. Multiple alternatively spliced variants, encoding the same protein, have been identified. [provided by RefSeq, Jul 2008]

**GeneCards Summary for GCNT1 Gene**

GCNT1 (Glucosaminyl (N-Acetyl) Transferase 1) is a Protein Coding gene. Among its related pathways are Metabolism and O-linked glycosylation. Gene Ontology (GO) annotations related to this gene include acetylglucosaminyltransferase activity and beta-1,3-galactosyl-O-glycosyl-glycoprotein beta-1,6-N-acetylglucosaminyltransferase activity. An important paralog of this gene is GCNT3.

**UniProtKB/Swiss-Prot Summary for GCNT1 Gene**

Glycosyltransferase that catalyzes the transfer of an N-acetylglucosamine moiety onto mucin-type core 1 O-glycan to form the branched mucin-type core 2 O-glycan. Mucin-type core 2 O-glycans play an important role in leukocyte extravasation as they serve as scaffolds for the display of the selectin ligand sialyl Lewis X by leukocytes.

**GSAP:**

**Entrez Gene Summary for GSAP Gene**

Accumulation of neurotoxic amyloid-beta is a major hallmark of Alzheimer disease (AD; MIM 104300). Formation of amyloid-beta is catalyzed by gamma-secretase (see PSEN1; MIM 104311), a protease with numerous substrates. PION, or GSAP, selectively increases amyloid-beta production through a mechanism involving its interaction with both gamma-secretase and its substrate, the amyloid-beta precursor protein (APP; MIM 104760) C-terminal fragment (APP-CTF) (He et al., 2010 [PubMed 20811458]).[supplied by OMIM, Nov 2010]

**GeneCards Summary for GSAP Gene**

GSAP (Gamma-Secretase Activating Protein) is a Protein Coding gene. Diseases associated with GSAP include Alzheimer Disease. Gene Ontology (GO) annotations related to this gene include amyloid-beta binding.

**UniProtKB/Swiss-Prot Summary for GSAP Gene**

Regulator of gamma-secretase activity, which specifically activates the production of amyloid-beta protein (amyloid-beta protein 40 and amyloid-beta protein 42), without affecting the cleavage of other gamma-secretase targets such has Notch. The gamma-secretase complex is an endoprotease complex that catalyzes the intramembrane cleavage of integral membrane proteins such as Notch receptors and APP (amyloid-beta precursor protein). Specifically promotes the gamma-cleavage of APP CTF-alpha (also named APP-CTF) by the gamma-secretase complex to generate amyloid-beta, while it reduces the epsilon-cleavage of APP CTF-alpha, leading to a low production of AICD.

**MEOX2:**

**Entrez Gene Summary for MEOX2 Gene**

 This gene encodes a member of a subfamily of non-clustered, diverged, antennapedia-like homeobox-containing genes. The encoded protein may play a role in the regulation of vertebrate limb myogenesis. Mutations in the related mouse protein may be associated with craniofacial and/or skeletal abnormalities, in addition to neurovascular dysfunction observed in Alzheimer's disease. [provided by RefSeq, Jul 2008]

**GeneCards Summary for MEOX2 Gene**

MEOX2 (Mesenchyme Homeobox 2) is a Protein Coding gene. Diseases associated with MEOX2 include Female Stress Incontinence and Low Compliance Bladder. Gene Ontology (GO) annotations related to this gene include DNA-binding transcription factor activity and RNA polymerase II proximal promoter sequence-specific DNA binding. An important paralog of this gene is MEOX1.

**UniProtKB/Swiss-Prot Summary for MEOX2 Gene**

Mesodermal transcription factor that plays a key role in somitogenesis and is required for sclerotome development (By similarity). Activates expression of CDKN1A and CDKN2A in endothelial cells, acting as a regulator of vascular cell proliferation. While it activates CDKN1A in a DNA-dependent manner, it activates CDKN2A in a DNA-independent manner (PubMed:22206000). May have a regulatory role when quiescent vascular smooth muscle cells reenter the cell cycle.

**MIER1:**

**Entrez Gene Summary for MIER1 Gene**

 This gene encodes a protein that was first identified in Xenopus laevis by its role in a mesoderm induction early response (MIER). The encoded protein functions as a transcriptional regulator. Alternatively spliced transcript variants encode multiple isoforms, some of which lack a C-terminal nuclear localization signal. [provided by RefSeq, May 2013]

**GeneCards Summary for MIER1 Gene**

MIER1 (MIER1 Transcriptional Regulator) is a Protein Coding gene. Diseases associated with MIER1 include Amelogenesis Imperfecta, Type Ig and Teeth Hard Tissue Disease. Gene Ontology (GO) annotations related to this gene include chromatin binding and obsolete signal transducer activity. An important paralog of this gene is MIER3.

**UniProtKB/Swiss-Prot Summary for MIER1 Gene**

Transcriptional repressor regulating the expression of a number of genes including SP1 target genes. Probably functions through recruitment of HDAC1 a histone deacetylase involved in chromatin silencing.

**USP38:**

**GeneCards Summary for USP38 Gene**

USP38 (Ubiquitin Specific Peptidase 38) is a Protein Coding gene. Among its related pathways are Ubiquitin-Proteasome Dependent Proteolysis and Integrated Breast Cancer Pathway. Gene Ontology (GO) annotations related to this gene include cysteine-type endopeptidase activity and thiol-dependent ubiquitinyl hydrolase activity. An important paralog of this gene is USP35.

**UniProtKB/Swiss-Prot Summary for USP38 Gene**

Deubiquitinating enzyme exhibiting a preference towards 'Lys-63'-linked ubiquitin chains.

**LINC0251:**

### No data available

**SPIN1:**

**GeneCards Summary for SPIN1 Gene**

SPIN1 (Spindlin 1) is a Protein Coding gene. Diseases associated with SPIN1 include Ovarian Cancer and Coproporphyria, Hereditary. Gene Ontology (GO) annotations related to this gene include methylated histone binding. An important paralog of this gene is SPIN3.

**UniProtKB/Swiss-Prot Summary for SPIN1 Gene**

Chromatin reader that specifically recognizes and binds histone H3 both trimethylated at 'Lys-4' and asymmetrically dimethylated at 'Arg-8' (H3K4me3 and H3R8me2a) and acts as an activator of Wnt signaling pathway downstream of PRMT2. In case of cancer, promotes cell cancer proliferation via activation of the Wnt signaling pathway (PubMed:24589551). Overexpression induces metaphase arrest and chromosomal instability. Localizes to active rDNA loci and promotes the expression of rRNA genes (PubMed:21960006). May play a role in cell-cycle regulation during the transition from gamete to embryo. Involved in oocyte meiotic resumption, a process that takes place before ovulation to resume meiosis of oocytes blocked in prophase I: may act by regulating maternal transcripts to control meiotic resumption.
