## Additional file 3 for "A systematic screen of breast cancer patients’ exomes for retrotransposon insertions reveals disease associated genes"

### Additional file 3: A short description as provided by “Entrez Gene Summary” (https://www.ncbi.nlm.nih.gov/gene), by “GeneCards Summary” (https://www.genecards.org) and by “UniProtKB/Swiss-Prot Summary” (https://uniprot.org) is given, if available, for each gene targeted by a retrotransposition event that could be confirmed by the presence of a TSD.

**GHR:**

### Entrez Gene Summary for GHR Gene

This gene encodes a member of the type I cytokine receptor family, which is a transmembrane receptor for growth hormone. Binding of growth hormone to the receptor leads to receptor dimerization and the activation of an intra- and intercellular signal transduction pathway leading to growth. Mutations in this gene have been associated with Laron syndrome, also known as the growth hormone insensitivity syndrome (GHIS), a disorder characterized by short stature. In humans and rabbits, but not rodents, growth hormone binding protein (GHBP) is generated by proteolytic cleavage of the extracellular ligand-binding domain from the mature growth hormone receptor protein. Multiple alternatively spliced transcript variants have been found for this gene. [provided by RefSeq, Jun 2011]

### GeneCards Summary for GHR Gene

### GHR (Growth Hormone Receptor) is a Protein Coding gene. Diseases associated with GHR include Laron Syndrome and Growth Hormone Insensitivity, Partial. Among its related pathways are Cytokine Signaling in Immune systemand Growth hormone receptor signaling. Gene Ontology (GO) annotations related to this gene include protein homodimerization activity and protein phosphatase binding. An important paralog of this gene is PRLR.

### UniProtKB/Swiss-Prot Summary for GHR Gene

### Receptor for pituitary gland growth hormone involved in regulating postnatal body growth. On ligand binding, couples to the JAK2/STAT5 pathway (By similarity).

### The soluble form (GHBP) acts as a reservoir of growth hormone in plasma and may be a modulator/inhibitor of GH signaling.

### Isoform 2 up-regulates the production of GHBP and acts as a negative inhibitor of GH signaling.

**GSTA5:**

**Entrez Gene Summary for GSTA5 Gene**

The glutathione S-transferases (GST; EC 2.5.1.18) catalyze the conjugation of reduced glutathiones and a variety of electrophiles, including many known carcinogens and mutagens. The cytosolic GSTs belong to a large superfamily, with members located on different chromosomes. For additional information on GSTs, see GSTA1 (MIM 138359).[supplied by OMIM, Sep 2008]

### GeneCards Summary for GSTA5 Gene

GSTA5 (Glutathione S-Transferase Alpha 5) is a Protein Coding gene. Among its related pathways are Glutathione metabolism and Platinum drug resistance. Gene Ontology (GO) annotations related to this gene include glutathione transferase activity. An important paralog of this gene is GSTA1.

**METTL3:**

### Entrez Gene Summary for METTL3 Gene

This gene encodes the 70 kDa subunit of MT-A which is part of N6-adenosine-methyltransferase. This enzyme is involved in the posttranscriptional methylation of internal adenosine residues in eukaryotic mRNAs, forming N6-methyladenosine. [provided by RefSeq, Jul 2008]

**GeneCards Summary for METTL3 Gene**

METTL3 (Methyltransferase Like 3) is a Protein Coding gene. Among its related pathways are Gene Expression and Circadian rythm related genes. Gene Ontology (GO) annotations related to this gene include RNA binding and mRNA (2'-O-methyladenosine-N6-)-methyltransferase activity.

**UniProtKB/Swiss-Prot Summary for METTL3 Gene**

The METTL3-METTL14 heterodimer forms a N6-methyltransferase complex that methylates adenosine residues at the N(6) position of some RNAs and regulates various processes such as the circadian clock, differentiation of embryonic and hematopoietic stem cells, cortical neurogenesis, response to DNA damage, differentiation of T-cells and primary miRNA processing (PubMed:22575960, PubMed:24284625, PubMed:25719671, PubMed:25799998, PubMed:26321680, PubMed:26593424, PubMed:27627798, PubMed:27373337, PubMed:27281194, PubMed:28297716, PubMed:30428350, PubMed:29506078, PubMed:29348140, PubMed:9409616). In the heterodimer formed with METTL14, METTL3 constitutes the catalytic core (PubMed:27627798, PubMed:27373337, PubMed:27281194). N6-methyladenosine (m6A), which takes place at the 5'-[AG]GAC-3' consensus sites of some mRNAs, plays a role in mRNA stability, processing, translation efficiency and editing (PubMed:22575960, PubMed:24284625, PubMed:25719671, PubMed:25799998, PubMed:26321680, PubMed:26593424, PubMed:28297716, PubMed:9409616). M6A acts as a key regulator of mRNA stability: methylation is completed upon the release of mRNA into the nucleoplasm and promotes mRNA destabilization and degradation (PubMed:28637692). In embryonic stem cells (ESCs), m6A methylation of mRNAs encoding key naive pluripotency-promoting transcripts results in transcript destabilization, promoting differentiation of ESCs (By similarity). M6A regulates the length of the circadian clock: acts as an early pace-setter in the circadian loop by putting mRNA production on a fast-track for facilitating nuclear processing, thereby providing an early point of control in setting the dynamics of the feedback loop (By similarity). M6A also regulates circadian regulation of hepatic lipid metabolism (PubMed:30428350). M6A regulates spermatogonial differentiation and meiosis and is essential for male fertility and spermatogenesis (By similarity). Involved in the response to DNA damage: in response to ultraviolet irradiation, METTL3 rapidly catalyzes the formation of m6A on poly(A) transcripts at DNA damage sites, leading to the recruitment of POLK to DNA damage sites (PubMed:28297716). M6A is also required for T-cell homeostasis and differentiation: m6A methylation of transcripts of SOCS family members (SOCS1, SOCS3 and CISH) in naive T-cells promotes mRNA destabilization and degradation, promoting T-cell differentiation (By similarity). Inhibits the type I interferon response by mediating m6A methylation of IFNB (PubMed:30559377). M6A also takes place in other RNA molecules, such as primary miRNA (pri-miRNAs) (PubMed:25799998). Mediates m6A methylation of Xist RNA, thereby participating in random X inactivation: m6A methylation of Xist leads to target YTHDC1 reader on Xist and promote transcription repression activity of Xist (PubMed:27602518). M6A also regulates cortical neurogenesis: m6A methylation of transcripts related to transcription factors, neural stem cells, the cell cycle and neuronal differentiation during brain development promotes their destabilization and decay, promoting differentiation of radial glial cells (By similarity). METTL3 mediates methylation of pri-miRNAs, marking them for recognition and processing by DGCR8 (PubMed:25799998). Acts as a positive regulator of mRNA translation independently of the methyltransferase activity: promotes translation by interacting with the translation initiation machinery in the cytoplasm (PubMed:27117702). Its overexpression in a number of cancer cells suggests that it may participate to cancer cell proliferation by promoting mRNA translation (PubMed:27117702).

**NEK10:**

### GeneCards Summary for NEK10 Gene

NEK10 (NIMA Related Kinase 10) is a Protein Coding gene. Gene Ontology (GO) annotations related to this gene include transferase activity, transferring phosphorus-containing groups and protein tyrosine kinase activity. An important paralog of this gene is NEK2.

**PTPN14:**

**Entrez Gene Summary for PTPN14 Gene**

The protein encoded by this gene is a member of the protein tyrosine phosphatase (PTP) family. PTPs are known to be signaling molecules that regulate a variety of cellular processes including cell growth, differentiation, mitotic cycle, and oncogenic transformation. This PTP contains an N-terminal noncatalytic domain similar to that of band 4.1 superfamily cytoskeleton-associated proteins, which suggested the membrane or cytoskeleton localization of this protein. It appears to regulate lymphatic development in mammals, and a loss of function mutation has been found in a kindred with a lymphedema-choanal atresia. [provided by RefSeq, Sep 2010]

### GeneCards Summary for PTPN14 Gene

PTPN14 (Protein Tyrosine Phosphatase Non-Receptor Type 14) is a Protein Coding gene. Diseases associated with PTPN14 include Choanal Atresia and Lymphedema and Lung Mucoepidermoid Carcinoma. Among its related pathways are Wnt / Hedgehog / Notch and PAK Pathway. Gene Ontology (GO) annotations related to this gene include phosphatase activity and transcription coregulator activity. An important paralog of this gene is PTPN21.

### UniProtKB/Swiss-Prot Summary for PTPN14 Gene

### Protein tyrosine phosphatase which may play a role in the regulation of lymphangiogenesis, cell-cell adhesion, cell-matrix adhesion, cell migration, cell growth and also regulates TGF-beta gene expression, thereby modulating epithelial-mesenchymal transition. Mediates beta-catenin dephosphorylation at adhesion junctions. Acts as a negative regulator of the oncogenic property of YAP, a downstream target of the hippo pathway, in a cell density-dependent manner. May function as a tumor suppressor.

**UPF2:**

**Entrez Gene Summary for UPF2 Gene**

This gene encodes a protein that is part of a post-splicing multiprotein complex involved in both mRNA nuclear export and mRNA surveillance. mRNA surveillance detects exported mRNAs with truncated open reading frames and initiates nonsense-mediated mRNA decay (NMD). When translation ends upstream from the last exon-exon junction, this triggers NMD to degrade mRNAs containing premature stop codons. This protein is located in the perinuclear area. It interacts with translation release factors and the proteins that are functional homologs of yeast Upf1p and Upf3p. Two splice variants have been found for this gene; both variants encode the same protein. [provided by RefSeq, Jul 2008]

### GeneCards Summary for UPF2 Gene

UPF2 (UPF2 Regulator of Nonsense Mediated MRNA Decay) is a Protein Coding gene. Among its related pathways are RNA transport and Viral mRNA Translation. Gene Ontology (GO) annotations related to this gene include RNA binding and binding.

### UniProtKB/Swiss-Prot Summary for UPF2 Gene

### Involved in nonsense-mediated decay (NMD) of mRNAs containing premature stop codons by associating with the nuclear exon junction complex (EJC). Recruited by UPF3B associated with the EJC core at the cytoplasmic side of the nuclear envelope and the subsequent formation of an UPF1-UPF2-UPF3 surveillance complex (including UPF1 bound to release factors at the stalled ribosome) is believed to activate NMD. In cooperation with UPF3B stimulates both ATPase and RNA helicase activities of UPF1. Binds spliced mRNA.

**ZNF442:**

### GeneCards Summary for ZNF442 Gene

ZNF442 (Zinc Finger Protein 442) is a Protein Coding gene. Among its related pathways are Herpes simplex virus 1 infection and Gene Expression. Gene Ontology (GO) annotations related to this gene include nucleic acid binding. An important paralog of this gene is ZNF443

**UniProtKB/Swiss-Prot Summary for ZNF442 Gene**

May be involved in transcriptional regulation.

**TMIGD3:**

### Entrez Gene Summary for TMIGD3 Gene

This gene encodes a transmembrane and immunoglobulin domain-containing protein. Alternative splicing results in multiple transcript variants, one of which shares its 5' terminal exon with that of the overlapping adenosine A3 receptor gene (GeneID:140), thus resulting in a fusion product. [provided by RefSeq, Nov 2014]

### GeneCards Summary for TMIGD3 Gene

TMIGD3 (Transmembrane and Immunoglobulin Domain Containing 3) is a Protein Coding gene. Gene Ontology (GO) annotations related to this gene include G protein-coupled receptor activity and G protein-coupled adenosine receptor activity. An important paralog of this gene is CD300E.

### UniProtKB/Swiss-Prot Summary for TMIGD3 Gene

### Isoform 1: Plays a suppressive role in osteosarcoma malignancy by inhibiting NF-kappa-B activity (PubMed:27886186).

**GPR42:**

### GeneCards Summary for GPR42 Gene

GPR42 (G Protein-Coupled Receptor 42 (Gene/Pseudogene)) is a Protein Coding gene. Among its related pathways are Peptide ligand-binding receptors. Gene Ontology (GO) annotations related to this gene include G protein-coupled receptor activity. An important paralog of this gene is FFAR3.

**UniProtKB/Swiss-Prot Summary for GPR42 Gene**

Probable G protein-coupled receptor that may be activated by a major product of dietary fiber digestion, the short chain fatty acids (SCFAs), and that may play a role in the regulation of whole-body energy homeostasis and/or in intestinal immunity.

**MIF-AS1:**

### GeneCards Summary for MIF-AS1 Gene

MIF-AS1 (MIF Antisense RNA 1) is an RNA Gene, and is affiliated with the non-coding RNA class. Diseases associated with MIF-AS1 include Nephrolithiasis, Calcium Oxalate.

**GSTT2B:**

**Entrez Gene Summary for GSTT2B Gene**

The protein encoded by this gene, glutathione S-transferase (GST) theta 2B (GSTT2B), is a member of a superfamily of proteins that catalyze the conjugation of reduced glutathione to a variety of electrophilic and hydrophobic compounds. Human GSTs can be divided into five main classes: alpha, mu, pi, theta, and zeta. The theta class includes GSTT1, GSTT2, and GSTT2B. GSTT2 and GSTT2B are nearly identical to each other, and share 55% amino acid identity with GSTT1. All three genes may play a role in human carcinogenesis. The GSTT2B gene is a pseudogene in some populations. [provided by RefSeq, Sep 2015]

### GeneCards Summary for GSTT2B Gene

GSTT2B (Glutathione S-Transferase Theta 2B (Gene/Pseudogene)) is a Protein Coding gene. Diseases associated with GSTT2B include Deafness, Autosomal Recessive 12. Among its related pathways are Glutathione metabolism and Platinum drug resistance. Gene Ontology (GO) annotations related to this gene include transferase activity and glutathione transferase activity. An important paralog of this gene is GSTT2.

### UniProtKB/Swiss-Prot Summary for GSTT2B Gene

### Conjugation of reduced glutathione to a wide number of exogenous and endogenous hydrophobic electrophiles. Has a sulfatase activity.

**CLVS1:**

### GeneCards Summary for CLVS1 Gene

CLVS1 (Clavesin 1) is a Protein Coding gene. Among its related pathways are Vesicle-mediated transport and Clathrin derived vesicle budding. Gene Ontology (GO) annotations related to this gene include transporter activity and phosphatidylinositol-3,5-bisphosphate binding. An important paralog of this gene is CLVS2.

**UniProtKB/Swiss-Prot Summary for CLVS1 Gene**

Required for normal morphology of late endosomes and/or lysosomes in neurons (By similarity). Binds phosphatidylinositol 3,5-bisphosphate (PtdIns(3,5)P2).

**DMBT1:**

**Entrez Gene Summary for DMBT1 Gene**

 Loss of sequences from human chromosome 10q has been associated with the progression of human cancers. This gene was originally isolated based on its deletion in a medulloblastoma cell line. This gene is expressed with transcripts of 6.0, 7.5, and 8.0 kb in fetal lung and with one transcript of 8.0 kb in adult lung, although the 7.5 kb transcript has not been characterized. The encoded protein precursor is a glycoprotein containing multiple scavenger receptor cysteine-rich (SRCR) domains separated by SRCR-interspersed domains (SID). Transcript variant 2 (8.0 kb) has been shown to bind surfactant protein D independently of carbohydrate recognition. This indicates that DMBT1 may not be a classical tumor suppressor gene, but rather play a role in the interaction of tumor cells and the immune system. [provided by RefSeq, Mar 2016]

### GeneCards Summary for DMBT1 Gene

DMBT1 (Deleted in Malignant Brain Tumors 1) is a Protein Coding gene. Diseases associated with DMBT1 include Dental Caries and Glioblastoma Multiforme. Among its related pathways are Salivary secretion and Surfactant metabolism. Gene Ontology (GO) annotations related to this gene include calcium-dependent protein binding and signaling pattern recognition receptor activity. An important paralog of this gene is CD163.

### UniProtKB/Swiss-Prot Summary for DMBT1 Gene

### May be considered as a candidate tumor suppressor gene for brain, lung, esophageal, gastric, and colorectal cancers. May play roles in mucosal defense system, cellular immune defense and epithelial differentiation. May play a role as an opsonin receptor for SFTPD and SPAR in macrophage tissues throughout the body, including epithelial cells lining the gastrointestinal tract. May play a role in liver regeneration. May be an important factor in fate decision and differentiation of transit-amplifying ductular (oval) cells within the hepatic lineage. Required for terminal differentiation of columnar epithelial cells during early embryogenesis. May function as a binding protein in saliva for the regulation of taste sensation. Binds to HIV-1 envelope protein and has been shown to both inhibit and facilitate viral transmission. Displays a broad calcium-dependent binding spectrum against both Gram-positive and Gram-negative bacteria, suggesting a role in defense against bacterial pathogens. Binds to a range of poly-sulfated and poly-phosphorylated ligands which may explain its broad bacterial-binding specificity. Inhibits cytoinvasion of S.enterica. Associates with the actin cytoskeleton and is involved in its remodeling during regulated exocytosis. Interacts with pancreatic zymogens in a pH-dependent manner and may act as a Golgi cargo receptor in the regulated secretory pathway of the pancreatic acinar cell.

**UBA6:**

**Entrez Gene Summary for UBA6 Gene**

Modification of proteins with ubiquitin (UBB; MIM 191339) or ubiquitin-like proteins controls many signaling networks and requires a ubiquitin-activating enzyme (E1), a ubiquitin conjugating enzyme (E2), and a ubiquitin protein ligase (E3). UBE1L2 is an E1 enzyme that initiates the activation and conjugation of ubiquitin-like proteins (Jin et al., 2007 [PubMed 17597759]).[supplied by OMIM, Mar 2008]

### GeneCards Summary for UBA6 Gene

UBA6 (Ubiquitin Like Modifier Activating Enzyme 6) is a Protein Coding gene. Among its related pathways are Protein ubiquitination and Ubiquitin mediated proteolysis. Gene Ontology (GO) annotations related to this gene include ligase activity and FAT10 activating enzyme activity. An important paralog of this gene is UBA1.

### UniProtKB/Swiss-Prot Summary for UBA6 Gene

### Activates ubiquitin by first adenylating its C-terminal glycine residue with ATP, and thereafter linking this residue to the side chain of a cysteine residue in E1, yielding a ubiquitin-E1 thioester and free AMP. Specific for ubiquitin, does not activate ubiquitin-like peptides. Differs from UBE1 in its specificity for substrate E2 charging. Does not charge cell cycle E2s, such as CDC34. Essential for embryonic development. Required for UBD/FAT10 conjugation. Isoform 2 may play a key role in ubiquitin system and may influence spermatogenesis and male fertility.

**C1orf194:**

### GeneCards Summary for C1orf194 Gene

C1orf194 (Chromosome 1 Open Reading Frame 194) is a Protein Coding gene. An important paralog of this gene is SLC49A3.

**PSMA7:**

**Entrez Gene Summary for PSMA7 Gene**

The 26S proteasome is a multicatalytic proteinase complex with a highly ordered structure composed of 2 complexes, a 20S core and a 19S regulator. The 20S core is composed of 4 rings of 28 non-identical subunits; 2 rings are composed of 7 alpha subunits and 2 rings are composed of 7 beta subunits. Proteasomes are distributed throughout eukaryotic cells at a high concentration and cleave peptides in an ATP/ubiquitin-dependent process in a non-lysosomal pathway. This gene encodes a member of the peptidase T1A family that functions as a 20S core alpha subunit. The encoded protein interacts with the hepatitis B virus X protein and plays a role in regulating hepatitis C virus internal ribosome entry site (IRES) activity, an activity essential for viral replication. The encoded protein also plays a role in the cellular stress response by regulating hypoxia-inducible factor-1alpha. A pseudogene of this gene is located on the long arm of chromosome 9. [provided by RefSeq, Jul 2012]

### GeneCards Summary for PSMA7 Gene

PSMA7 (Proteasome Subunit Alpha 7) is a Protein Coding gene. Diseases associated with PSMA7 include Plethora Of Newborn and Hypoxia. Among its related pathways are Cellular Senescence (REACTOME) and TCR signaling (REACTOME). Gene Ontology (GO) annotations related to this gene include identical protein binding and threonine-type endopeptidase activity. An important paralog of this gene is PSMA8.

### UniProtKB/Swiss-Prot Summary for PSMA7 Gene

Component of the 20S core proteasome complex involved in the proteolytic degradation of most intracellular proteins. This complex plays numerous essential roles within the cell by associating with different regulatory particles. Associated with two 19S regulatory particles, forms the 26S proteasome and thus participates in the ATP-dependent degradation of ubiquitinated proteins. The 26S proteasome plays a key role in the maintenance of protein homeostasis by removing misfolded or damaged proteins that could impair cellular functions, and by removing proteins whose functions are no longer required. Associated with the PA200 or PA28, the 20S proteasome mediates ubiquitin-independent protein degradation. This type of proteolysis is required in several pathways including spermatogenesis (20S-PA200 complex) or generation of a subset of MHC class I-presented antigenic peptides (20S-PA28 complex). Inhibits the transactivation function of HIF-1A under both normoxic and hypoxia-mimicking conditions. The interaction with EMAP2 increases the proteasome-mediated HIF-1A degradation under the hypoxic conditions. Plays a role in hepatitis C virus internal ribosome entry site-mediated translation. Mediates nuclear translocation of the androgen receptor (AR) and thereby enhances androgen-mediated transactivation. Promotes MAVS degradation and thereby negatively regulates MAVS-mediated innate immune response.

**UQCR10:**

**Entrez Gene Summary for UQCR10 Gene**

UCRC is a subunit of mitochondrial complex III (ubiquinol-cytochrome c reductase; EC 1.10.2.2), which forms the middle segment of the respiratory chain of the inner mitochondrial membrane (Schagger et al., 1995 [PubMed 8592474]).[supplied by OMIM, Mar 2008]

### GeneCards Summary for UQCR10 Gene

UQCR10 (Ubiquinol-Cytochrome C Reductase, Complex III Subunit X) is a Protein Coding gene. Among its related pathways are Respiratory electron transport, ATP synthesis by chemiosmotic coupling, and heat production by uncoupling proteins. and Metabolism. Gene Ontology (GO) annotations related to this gene include ubiquinol-cytochrome-c reductase activity.

### UniProtKB/Swiss-Prot Summary for UQCR10 Gene

### This is a component of the ubiquinol-cytochrome c reductase complex (complex III or cytochrome b-c1 complex), which is part of the mitochondrial respiratory chain. This subunit interacts with cytochrome c1 (By similarity).

**MRPL50:**

**Entrez Gene Summary for MRPL50 Gene**

Mammalian mitochondrial ribosomal proteins are encoded by nuclear genes and help in protein synthesis within the mitochondrion. Mitochondrial ribosomes (mitoribosomes) consist of a small 28S subunit and a large 39S subunit. They have an estimated 75% protein to rRNA composition compared to prokaryotic ribosomes, where this ratio is reversed. Another difference between mammalian mitoribosomes and prokaryotic ribosomes is that the latter contain a 5S rRNA. Among different species, the proteins comprising the mitoribosome differ greatly in sequence, and sometimes in biochemical properties, which prevents easy recognition by sequence homology. This gene encodes a putative 39S subunit protein and belongs to the L47P ribosomal protein family. Pseudogenes corresponding to this gene are found on chromosomes 2p, 2q, 5p, and 10q. [provided by RefSeq, Jul 2008]

### GeneCards Summary for MRPL50 Gene

MRPL50 (Mitochondrial Ribosomal Protein L50) is a Protein Coding gene. Among its related pathways are Mitochondrial translation and Organelle biogenesis and maintenance.
